# A Polygenic Route to Thermal Melanism and high-elevation adaptation in Honey Bees

**DOI:** 10.64898/2026.05.18.725832

**Authors:** Marco Mazzoni, Florian Loiolt, Sonja Kersten, Mark Otieno, Henry Njovu, Julius Vincent Lasway, Ricarda Scheiner, Martin Hasselmann

## Abstract

Thermal melanism, whereby darker pigmentation occurs in colder environments, is a widespread adaptive pattern, yet its genetic and physiological basis in eusocial insects remains poorly understood. East African honey bees (Apis mellifera) occupy steep elevational gradients in which highland populations are darker, larger, and more cold tolerant than lowland conspecifics. Here, we integrated population genomics, structural variation, reciprocal translocations, and functional genomics to dissect the basis of elevational melanism in honey bees. 139 workers from five East African mountain systems were sampled and phenotyped, and whole-genome resequencing was performed. A genome-wide association study identified 977 SNPs significantly associated with abdominal pigmentation, with a single pronounced peak at the ebony ortholog AmEbony on chromosome CM009931.2. Allelic variation at AmEbony formed three genetic clusters that closely tracked a continuous pigmentation gradient, and 77 highly divergent SNPs (DXY = 1) almost perfectly discriminated dark from light bees. Two large inversions (r7, r9), previously linked to high-elevation adaptation, were detected across additional mountain systems and were enriched in dark, highland genomic backgrounds, but did not replace AmEbony as the primary pigmentation locus. Reciprocal highland–lowland translocations revealed lineage-specific yet convergent expression shifts along detoxification/immune, chemosensory, proteostasis, and cuticle axes. CRISPR/Cas9-mediated disruption of AmEbony in A. m. carnica altered pigmentation and induced coherent changes in head transcriptomes, notably in odorant-binding and redox-related modules. Our results demonstrate that thermal melanism in a eusocial pollinator is governed by a top-heavy polygenic architecture centered on AmEbony, linking mechanistically naturally segregating pigmentation alleles to gene regulatory reprogramming and high-elevation adaptation.

## Introduction

Adaptation to heterogeneous environments is a central driver of evolutionary divergence and speciation. In ectotherms, two key traits, body pigmentation and body size, are critical for coping with the thermal and physiological challenges imposed by elevational gradients. The Thermal Melanism Hypothesis (TMH) proposes that darker pigmentation enhances solar-heat absorption and thus confers a selective advantage in cooler, high-altitude environments(Clusella Trullas et al. 2007; Clusella - Trullas et al. 2008). In parallel, larger body size can increase thermal inertia and improve performance under low-temperature conditions (Stevenson 1985). Empirical support for TMH has accumulated across a broad range of insect orders. Among Lepidoptera, Pieridae and Nymphalidae species exhibit increasingly dark wing patterns with increasing altitude (Kingsolver 1985; Markl et al. 2022), a pattern corroborated by large-scale surveys of North American butterflies across Lycaenidae, Pieridae, Nymphalidae and Papilionidae (Stelbrink et al. 2019). Comparable elevational and seasonal clines in pigmentation have been reported in Odonata (Novella-Fernandez et al. 2023), Orthoptera (Harris et al. 2013), Diptera (Freoa et al. 2023), and Hymenoptera, including paper wasps (de Souza et al. 2020). Analogous patterns are evident in vertebrates: color polymorphism in the rock pocket mouse (Nachman et al. 2003; Hoekstra et al. 2004) and Peromyscus mice (Barrett et al. 2019; Hager et al. 2022) are tightly associated with environmental heterogeneity and local adaptation. Together, these studies highlight thermal melanism as a widespread and powerful mechanism facilitating adaptation to high-altitude and other thermally challenging environments.

The western honey bee, *Apis mellifera*, provides an exceptional model for dissecting polygenic adaptation across climatic gradients (Wallberg et al. 2014). In East African mountain systems, two subspecies, *A. m. monticola* and *A. m. scutellata* co-occur but are ecologically segregated: *A.m. monticola* occupies rather cool, humid montane forests, while *A.m. scutellata* dominates hot, arid savannahs (Ruttner 1988). Highland bees exhibit several distinguishing characteristics. They are darker in color, larger in size, and capable of flight at lower temperatures (Ruttner 1988; Gruber et al. 2013; Wallberg et al. 2017), consistent with TMH. Yet, population genomic analyses of Kenyan honey bee revealed that *A.m. monticola* and *A.m. scutellata* share approximately 98.5% of their genome whereas the remaining fraction showing pronounced differentiation (F_ST_ 0.6-0.8), driven by chromosomal inversions (r7, r9) (Wallberg et al. 2017; Christmas et al. 2018). Recent work has confirmed these inversions in Ugandan honey bee populations and linked them to elevation related gene expression differences, particularly in stress tolerance and cuticular hydrocarbon pathways (Mazzoni et al. 2025). However, these inversions do not fully explain the observed variation in pigmentation. Instead, genome-wide selection scans and association mapping in Kenyan populations have pinpointed two pigmentation genes, orthologous to *Drosophila melanogaster tan* and *ebony* (Wittkopp et al. 2002; True et al. 2005; Massey and Wittkopp 2016), that show strong correlation with elevational habitat (Mazzoni et al. 2025).

Establishing causal links between specific genes and adaptive phenotypes requires functional validation (Barrett and Hoekstra 2011; Stern 2013). CRISPR/Cas9 genome editing has become a powerful approach for dissecting gene function in diverse organisms (Jinek et al. 2012). In Drosophila, *tan* and *ebony* knockouts alter both, melanin deposition and cuticular hydrocarbon profiles (Massey et al. 2019). In honey bees, targeted disruption of *yellow* (y) affects expression of key melanogenesis enzymes such as laccase 2 and aaNAT, with consequences for pigmentation and biogenic amine metabolism (Arakane et al. 2005; Nie et al. 2021). Despite these advances, a mechanistic, experimentally validated link between naturally segregating pigmentation alleles and elevational adaptation in a eusocial insect remains lacking.

Here, we integrate population genomic analyses, transcriptome analyses and functional genomics to elucidate the genetic architecture and regulatory basis of high-altitude pigmentation in East African honey bees. We tested the hypothesis that elevational adaptation in these populations is driven by a polygenic architecture linked to environmental conditions. We assembled and phenotyped population samples from multiple African mountain systems, the Rwenzori Mountains (Uganda), Mt. Kilimanjaro and Mt. Meru (Tanzania), and Mt. Kenya and Mt. Elgon (Kenya), and combined these data with genomic and color phenotypes from (Wallberg et al. 2017). A genome-wide association study (GWAS) reveals a polygenic basis of elevational melanism, implicating *ebony*, *tan*, and additional loci beyond the major inversion regions. To directly connect environmental variation with gene regulatory responses, we conducted a reciprocal transplant experiment of highland and lowland colonies and quantified their transcriptional response to novel thermal environments. Furthermore, to establish causality at a key candidate locus, we used CRISPR/Cas9 to generate *ebony* knockouts in *A. m. carnica* and performed tissue-specific RNA-seq on head, thorax, and abdomen. This dual approach, linking landscape-level allele frequency patterns and expression shifts with precise functional perturbation, demonstrates how a gene perturbation reshapes gene regulatory networks and confirms a causal role of *AmEbony* in honey bee pigmentation.

Collectively, our study (i) provides the first integrative, multi-scale demonstration that thermal melanism in a eusocial pollinator has a polygenic basis rooted in canonical insect pigmentation genes, (ii) shows that major chromosomal inversions alone cannot account for elevational pigmentation clines in East African honey bees, and (iii) establishes a direct functional link between a naturally segregating ebony allele, transcriptional reprogramming, and an adaptive pigmentation phenotype. This work advances mechanistic understanding of thermal melanism and offers a general framework for connecting population genomic signals of selection to experimentally validated causal loci in an ecologically important species.

## Results

### Identifying genomic regions linked to pigmentation differentiation

We analyzed 139 honey bees sampled across five major East Africa mountain systems that span pronounced elevational gradients: Mount Elgon in western Kenya (maximum elevation ∼4,321 m; maximum sample elevation 2600m), the Rwenzori Mountain system in Uganda (maximum elevation ∼5,109 m: maximum sample elevation: ∼2700m), Mount Kenya in central Kenya (maximum elevation ∼5,199 m; maximum sample elevation: ∼2800m), Mount Kilimanjaro in northern Tanzania (maximum elevation ∼5,895 m), and Mount Meru in northern Tanzania (maximum elevation ∼4,566 m; maximum sample elevation: ∼2800m; Supplementary Table 1). These systems were selected because they encompass steep climatic gradients over short geographic distances and harbor natural populations of *A. mellifera* adapted to contrasting thermal and ecological conditions.

Each individual was assigned a quantitative abdominal pigmentation score (Supplementary Figure 1), derived from standardized, high-resolution digital images obtained under controlled laboratory conditions, thereby minimizing phenotyping bias. Consistent with our a priori expectation of thermal melanism, abdominal pigmentation increased markedly with elevation (Figure 1A), with high-altitude bees exhibiting markedly darker abdomens than their lowland counterparts.

**Figure 1.**
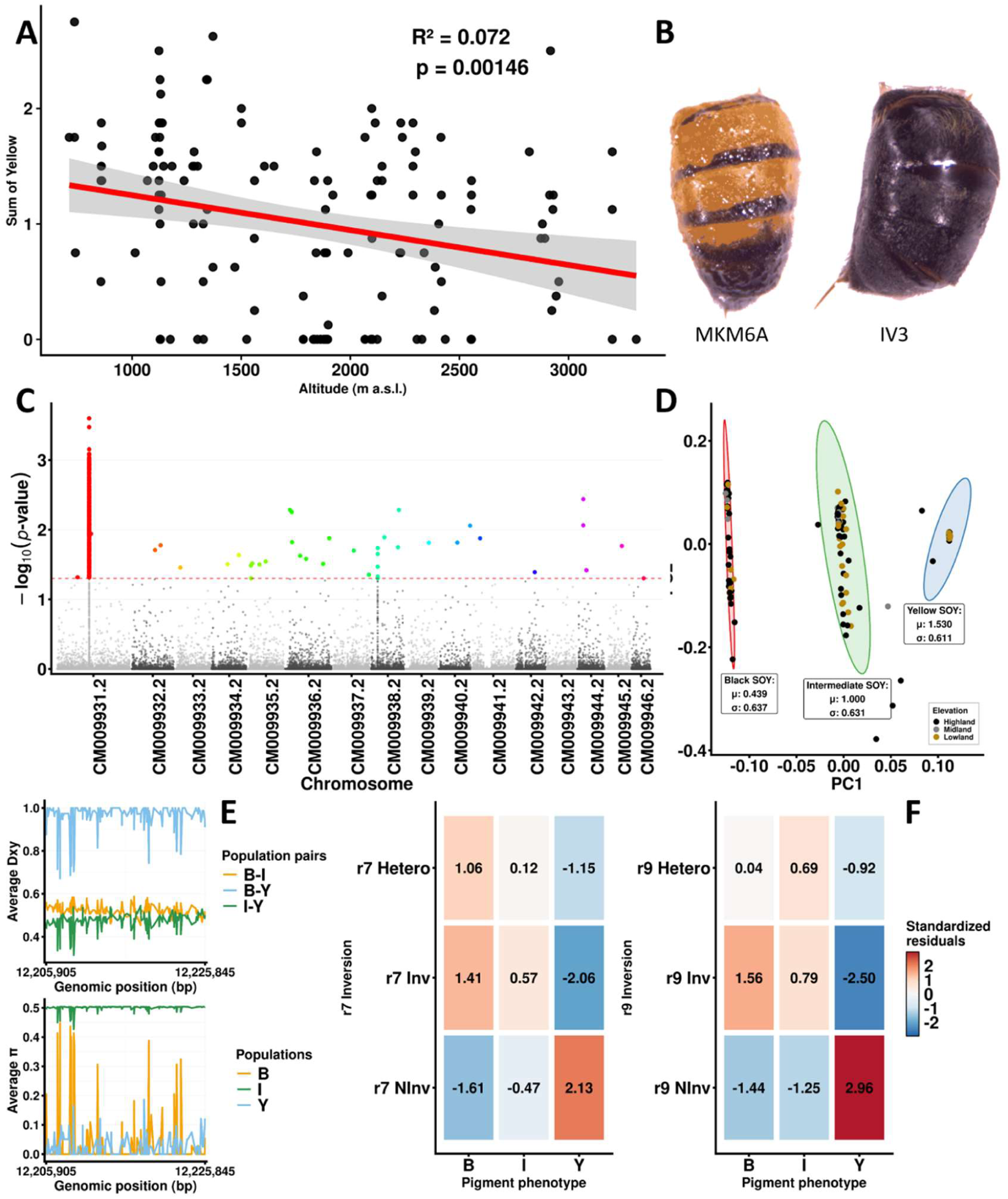
Altitudinal variation in pigmentation and associated genomic signals. (A) Relationship between altitude (m a.s.l.) and the pigmentation index “Sum of Yellow” (SOY): the yellow component decreases with increasing altitude (red regression line; grey band = confidence interval). R² and p value are reported. (B) Representative individuals from the two extreme phenotype clusters (yellow vs black abdomen). (C) Genome-wide association (Manhattan) plot for the pigmentation phenotype: y-axis shows −log10(p); the dashed line indicates the significance threshold and highlighted points denote the strongest association signals. (D) PCA-based clustering of individuals using SOY-related metrics, showing three groups (black - B, intermediate -I, yellow -Y ellipses) and their altitudinal origin; insets report group summary statistics (mean μ and standard deviation σ). (E) Population-genetic statistics across the candidate interval: mean divergence (Dxy) between phenotype-group pairs (B–I, B–Y, I–Y) and nucleotide diversity (π) within each group along genomic coordinates. (F) Association between pigmentation phenotype (B/I/Y) and inversion genotypes (heterozygous, inversion homozygous, non-inversion homozygous) for r7 and r9: heatmaps show standardized residuals (red = enrichment relative to expectation; blue = depletion).

Whole-genome sequencing of the 139 individuals initially identified approximately 10 million SNPs. After stringent quality filtering based on read coverage, minor allele frequency, and missing data (see Material and Methods), we retained 6,631,751 high-quality SNPs for downstream analyses. Because abdominal pigmentation could not be scored for one individual, the final association analyses were conducted on 138 phenotyped bees (Supplementary Table 2). This curated SNP set provided the basis for all subsequent genome-wide association and population genomic analyses.

We performed a genome-wide association study (GWAS) using a linear mixed model implemented in GEMMA, incorporating all 6,631,751 SNPs and a single covariate to account for potential confounding factors. The model estimated a proportion of phenotypic variance explained (PVE) of 0.9999 ± 0.0067, indicating that nearly the entirety of phenotypic variation in abdominal pigmentation was attributable to genetic factors captured by the model. The estimates of genetic variance (v_g_ = 1.8885) and negligible residual variance (v_e_ ≈ 1.9 × 10^−5^) in the null model further underscored the substantial genetic component underlying the trait. The restricted maximum-likelihood (REML) log-likelihood of the null model (–90.89) and the maximum-likelihood estimate (–86.36) indicate a well-fitted baseline model a trait under strong genetic control.

Association statistics were corrected for multiple testing using the Benjamini–Hochberg (BH) false discovery rate (FDR) procedure (adjusted p < 0.05). We identified 977 SNPs significantly associated with abdominal pigmentation, the overwhelming majority of which were concentrated within a discrete region on chromosome CM009931.2. The Manhattan plot (Figure 1B) revealed a single, pronounced association peak spanning positions 12,205,905 to 12,225,845 bp, with a dense duster of significant variants. Concordant peaks in absolute divergence (D_XY_) in this interval supports a tightly linked haplotype, indicative for a major locus underlying abdominal pigmentation. A limited number of additional loci located on other chromosomes exhibited moderate association signals but did not surpass the genome-wide significance threshold.

The zoomed-in view of the Manhattan plot (Supplementary Figure 2) further resolves the lead SNPs within this region, revealing a dense cluster of strongly associated variants. Together, these patterns indicate that the altitudinally associated variation in abdominal pigmentation among East African honey bees is dominated by a single genomic locus, with relatively minor contributions from the remainder of the genome. This contrasts with the largely polygenic architecture expected for many quantitative traits and highlights a major-effect locus driving elevational melanism in this system.

To reduce proximal contamination and verify that the observed association peak was not an artefact of the mixed-model specification, we additionally implemented a leave-one-chromosome-out (LOCO) linear mixed model. In this framework, when testing SNPs on chromosome i, the genomic relationship matrix (GRM) is re-estimated excluding all variants on chromosome i, so that local effects are not absorbed into the random effect term. Re-running the GWAS under the LOCO scheme with the same set of individuals, genotypes, and covariates yielded a concordant major peak on CM009931.2, confirming the robustness of the signal (Supplementary Figure 3 – 4).

Within this associated region, several annotated gene models are present. Reciprocal BLASTP searches between *Drosophila melanogaster ebony* (FBgn0000527) and the *Apis mellifera* candidate LOC409109 yielded concordant top hits in both directions (E = 0.0; bit score = 798; 48% identity over 883 aligned amino acids; 67% positives; 3% gaps), strongly supporting LOC409109 as the *A. mellifera* ortholog of *ebony*. To further validate orthology and rule out paralogy, we reconstructed a phylogeny of *ebony* homologs across hymenoptera order. The *A. mellifera* sequence clustered with apoidea e*bony* homologs and showed, as expected, a high evolutionary divergence to *D. melanogaster* (Supplementary Figure 5). For clarity of presentation, we refer to this locus as “*AmEbony*” throughout the manuscript.

To characterize the genetic structure at *AmEbony*, we extracted all GWAS-significant SNPs located within this locus and summarized their allelic variation using a principal component analysis (PCA). This analysis revealed three genetically distinct clusters (Figure 1D) that mapped closely onto the observed abdominal pigmentation phenotypes. One cluster comprised predominantly darkly pigmented (black) bees with an average pigmentation score of 0.439 ± 0.637, a second cluster comprised individuals with intermediate pigmentation with an average score of 1.00 ± 0.631, and a third cluster included predominantly lightly pigmented (yellow) bees with an average score of 1.53 ± 0.611. The concordance between the PCA-derived clusters and phenotypic pigmentation scores underscores the central role of the *AmEbony* locus in shaping the observed variation in abdominal melanism.

To investigate the potential functional relevance of the associated variants, we annotated all significant SNPs within *AmEbony* using SnpEff (Cingolani et al. 2012). Although most significant SNPs were intronic, a subset was predicted to affect regulatory or coding regions. Specifically, we identified 3 SNPs located in the 3’ untranslated region (3’UTR), 13 SNPs in the 5’ untranslated region (5’UTR), 13 missense substitutions, and 26 synonymous substitutions within the coding sequence. This composition suggests that variation at *AmEbony* likely contributes to pigmentation differences via a combination of *cis*-regulatory changes and amino-acid substitutions, in addition to extensive intronic variation that may affect splicing or other regulatory processes.

To identify the variants most strongly associated with pigmentation divergence among the three phenotypic groups, we calculated standard population genomic statistics using pixy (Korunes and Samuk 2021) (Figure 1E), including F_ST_, D_XY_, and nucleotide diversity (π), across the pigmentation-defined clusters (yellow, intermediate, and black). As expected, the general patterns observed for F_ST_ and D_XY_ were largely congruent. For downstream analyses, we focused on D_XY_, as it measures absolute genetic divergence between populations and is less sensitive to differences in within-population diversity. Using this approach, we identified 77 SNPs within *AmEbony* with D_XY_ = 1 between the predominantly black cluster and the other pigmentation groups. Among these highly divergent variants, 2 SNPs were located within the 5’UTR, 5 were synonymous, 5 were missense substitutions, and the remaining SNPs resided in intronic regions (Supplementary Table 3).

Strikingly, these 77 SNPs almost completely discriminate black-pigmented bees from yellow-pigmented bees. Their allele frequencies shift gradually across the pigmentation spectrum: the alleles associated with dark pigmentation are rare in the yellow cluster, intermediate in frequency in the phenotypically intermediate cluster, and approach fixation in the black cluster. This allele-frequency cline mirrors the continuous phenotypic gradient in abdominal pigmentation and highlights strong, phenotype-correlated differentiation at AmEbony.

The concordance between allele frequency trajectories and pigmentation scores further reinforces the pivotal contribution of these SNPs predominantly intronic, with a subset affecting regulatory or coding regions to the altitudinally associated variation in abdominal melanism.

### Inversions associated with high-elevation adaptation are also connected to melanism

Previous genomic studies of East African honey bees identified two large chromosomal inversions - hereafter r7 and r9 - on chromosome 7 (CM009937.2) and chromosome 9 (CM009939.2), respectively, that are associated with local adaptation to high-elevation environments (Wallberg et al. 2014; Wallberg et al. 2017; Christmas et al. 2018; Mazzoni et al. 2025). In addition, a region of elevated genetic differentiation between highland and lowland populations has been reported at the proximal end of chromosome 5 (CM009935.2). Although this region shows patterns suggestive of an inversion, its structural status remains unresolved in the absence of long-read data; we therefore refer to it as the r5 region (Mazzoni et al. 2025). Our sampling included bees from Mount Kilimanjaro and Mount Meru in northern Tanzania, two high-elevation regions where the presence of r7 and r9 had been hypothesized based on ecological and altitudinal patterns but not directly demonstrated (Wallberg et al. 2017). Genotyping of these Tanzanian samples confirmed the presence of both r7 and r9, providing the first formal documentation of these inversions in Kilimanjaro and Meru populations. This extends the known geographic range of these structural variants beyond previously characterized East African mountain systems (Supplementary Figure 6; Supplementary Figure 7).

To examine how these genomic features relate to the pigmentation clusters defined by the *AmEbony-*associated SNPs, we assigned each of the 138 phenotyped bees to one of the three pigmentation clusters (black, intermediate, yellow) and, independently assigned inversion genotypes for r7 and r9 (homozygous for the reference orientation, heterozygous, or homozygous for the inverted orientation) based on diagnostic SNPs (Wallberg et al. 2017; Christmas et al. 2018; Mazzoni et al. 2025). For the r5 region, individuals were characterized as carrying highland-like or lowland-like haplotypes based on previously reported diagnostic SNPs. We then compared the distribution of inversion genotypes across the pigmentation-defined clusters. This analysis revealed a clear, non-random association between the darker abdominal pigmentation and the derived inverted orientations of both r7 and r9. Chi-square tests rejected independence between pigmentation cluster and inversion genotype for r7 ( χ² = 16.33, p = 0.0026) and r9 (χ² = 23.02, p = 0.0001) (Figure 1F, full contingency tables in Supplementary Table 4). In contrast, the r5 region showed no such pattern: haplotype frequencies did not differ significantly among the pigmentation clusters (χ² = 6.47,p = 0.167), despite pronounced highland–lowland differentiation at this locus (Mazzoni et al. 2025).

Taken together, these results show that the two previously characterized inversions, r7 and r9, which are strongly associated with high-elevation adaptation, are also enriched in the pigmentation cluster characterized by darker abdominal coloration. This pattern is consistent with a shared geographic and ecological context: high, cool environments favoring both inverted haplotypes and dark pigmentation rather than a direct causal effect of the inversions on the pigmentation phenotype itself. In contrast, the r5 region, despite its strong altitudinal differentiation, shows no analogous concordance with pigmentation clusters. These findings reinforce that *AmEbony* represents the primary locus directly underlying abdominal melanism, whereas r7 and r9 mark broader highland genomic backgrounds pointing to relevant genetic component therein for high-elevation adaptation.

### CRISPR-Cas9-induced *AmEbony* disruption is consistent with pigmentation change and chemosensory transcript shifts

To functionally test the role of *AmEbony* in abdominal pigmentation, we employed CRISPR/Cas9-mediated gene disruption. Guide RNAs were designed to target conserved exonic regions of *AmEbony*, with the aim of generating frameshift mutations predicted to abolish gene function. Ribonucleoprotein complexes of Cas9 and specific sgRNA were microinjected into one-cell stage honey bee embryos to induce targeted mutagenesis. Because genome editing is not yet feasible in the East African populations analyzed in our GWAS due to a lack of infrastructure, we conducted the experimental work in *Apis mellifera carnica*, a widely used experimental subspecies that provides a tractable model for functional genetics. This design assumes conservation of the core melanization pathway across *A. mellifera* subspecies while acknowledging that regulatory context and standing genetic variation may differ.

Edited individuals were genotyped using fluorescence-labeled amplicon (FLA) analysis, enabling unambiguous discrimination between wild-type (WT) and knockout (KO) genotypes (Figure 2A). The CRISPR-mediated disruption of *AmEbony* produced indel mutations at the targeted exons sites, consistent with loss-of-function mutations in the encoded protein (Figure 2B). We did not quantify somatic mosaicism or off-target edits at the tissue level, because we injected embryos at the one-cell stage. We successfully established both WT and *AmEbony* KO individuals within laboratory colonies (Supplementary Figure 8), providing a controlled framework to assess the direct phenotypic consequences of *AmEbony* disruption on abdominal pigmentation and to characterize associated changes in gene expression.

**Figure 2.**
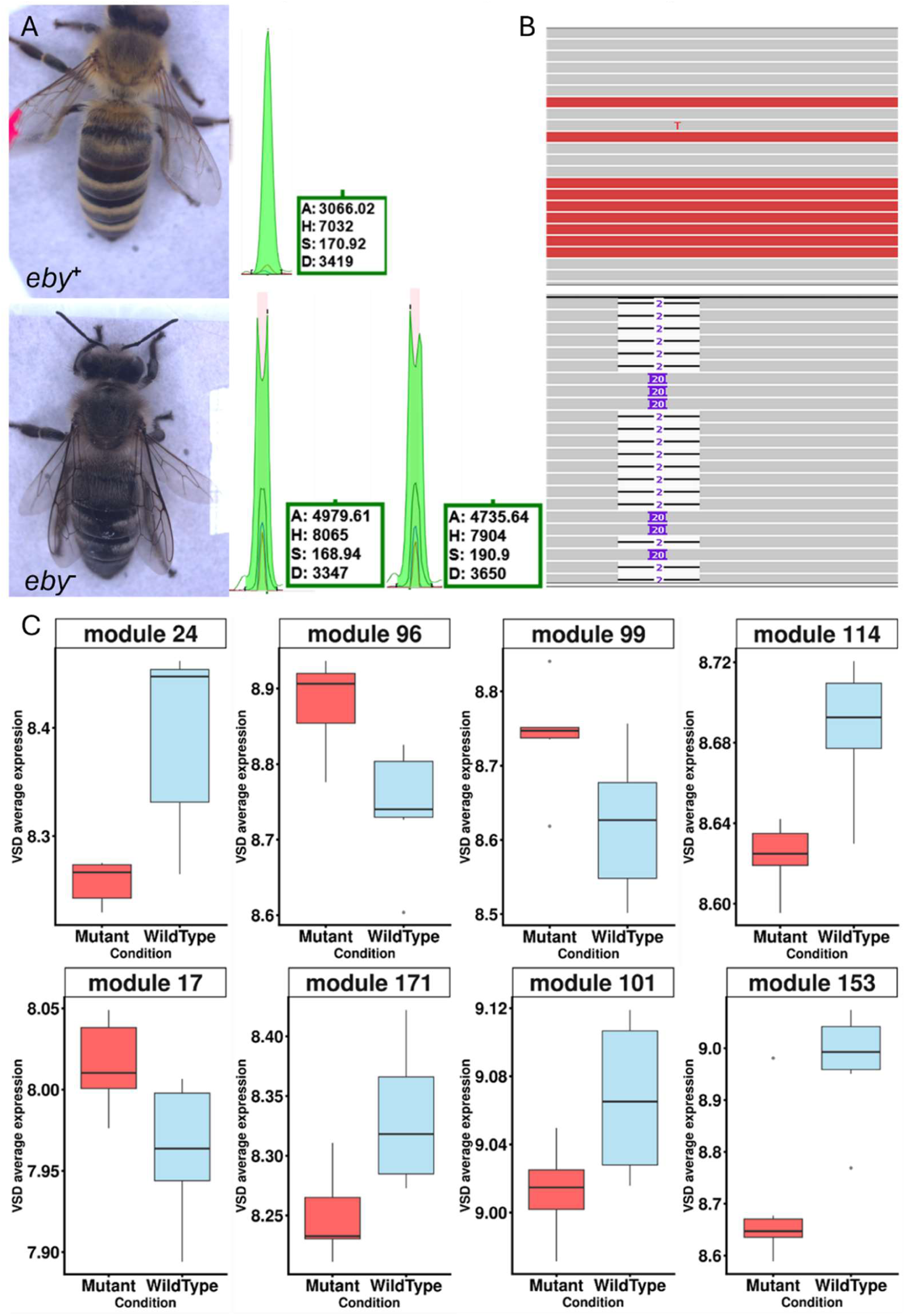
CRISPR/Cas9 validation of *ebony* editing and GENIE3-based transcriptomic module differences. (A) Representative phenotypes of a wild-type individual (eby^+^, yellow pigmentation) and a CRISPR-edited knockout (eby^−^, dark pigmentation). Fragment length analysis (FLA) of the targeted locus (green electropherogram peaks) supports successful genome editing, with edited individuals showing fragment patterns consistent with indel formation at the CRISPR cut site. (B) Read alignments across the targeted region (IGV-style view) further confirm editing at the expected locus, with recurrent indel classes (e.g., 2 bp and 20 bp events, indicated in purple). (C) Gene regulatory network inference using GENIE3 (run 1) identified eight expression modules significantly associated with genotype (significance assessed within the GENIE3-based analysis workflow; p-values reported in Methods). Boxplots show mean (+/- standard deviation) variance-stabilized expression (VSD average expression) for each significant module (24, 96, 99, 114, 17, 171, 101, 153) in mutant (red) versus wild-type (blue) samples.

### Expression patterns of the head-derived co-expression modules

To explore how *AmEbony* loss-of-function reshapes regulatory networks, we inferred co-expression modules from head transcriptomes of WT and *AmEbony* KO bees using GENIE3. This analysis yielded ∼175 modules, capturing structured transcriptional responses to *AmEbony* disruption.

GENIE3 relies on tree-based ensemble regressors (Random Forests/Extra-Trees). Therefore, some stochasticity in edge inference was expected. To evaluate the robustness of the inferred functional signals, we repeated the full GENIE3 analysis five times using different random seeds and increased the number of trees from 1,000 to 2,000 per run. We then performed GO enrichment analyses for each run and summarized enrichment reproducibility at the level of GO-term presence/absence per run. Across runs (assessing enrichment reproducibility at the level of “presence in a run”, i.e., detected in at least one model within that run), we observed 102 unique enriched GO terms. Only one GO term, oxidoreductase activity (GO:0016491),was recovered in all 5/5 runs, and only one, odorant binding (GO:0005549), was recovered in 4/5 runs (Supplementary Table 5). Eleven GO terms were supported in 3/5 runs, whereas the majority (79/102) appeared in only a single run. Based on this empirical stability assessment, we consider enrichments supported by ≥4/5 runs as the most robust functional signals, and GO terms observed in 3/5 runs as suggestive patterns that should be interpreted cautiously and preferably in conjunction with convergent evidence from gene-level analyses or independent datasets.

The WT-biased modules were enriched for mitochondrial and energy metabolism pathways, including oxidative phosphorylation, components of the respiratory electron-transport chain, ATP metabolic processes, and mitochondrial complex assembly. In contrast, the KO-biased modules were enriched for redox- and antioxidant-related functions, with oxidoreductase activity (GO:0016491) representing the single most reproducible enrichment across all runs (5/5; Supplementary Table 5). Notably, the head differentially expressed genes (DEGs) were strongly enriched for odorant-binding proteins, and odorant binding (GO:0005549) was among the most reproducible (4/5 runs; Supplementary Table 5). This indicates that *AmEbony* disruption in the head affects not only core metabolic and redox-related processes but also chemosensory pathways, particularly those involving odorant-binding proteins.

Together, these results show that loss of *AmEbony* triggers a modular and functionally coherent transcriptional response in head tissue, characterized by shifts in mitochondrial energy metabolism, redox balance, and odorant-binding capacity. At the same time, the run-to-run variability in many specific GO terms underscores the importance of replication and stability assessment in network-based enrichment analyses. In combination with the strong GWAS signal and the inversion–melanism associations, these transcriptomic changes provide functional support for a central role of *AmEbony* in linking pigmentation, physiology, and chemosensory function in honey bees.

### Reciprocal translocation experiments reveal lineage-specific yet convergent expression shifts along coherent physiological axes

Reciprocal translocation of colonies across elevations shifted gene expression along distinct, physiologically coherent axes that depended on lineage of origin but tended to converge toward the transcriptional profile characteristic of the destination habitat. For each contrast, the non-translocated group served as the baseline (LL for LLvsLH; HH for HHvsHL; LL for HHvsLL). In highland-origin colonies moved downslope (HH vs HL, baseline HH), we identified 58 DEGs (36 higher in HH; 22 higher in HL; median |log2FC| 1.47). Genes upregulated in HL were significantly enriched for monooxygenase activity and heme/iron binding and included multiple cytochrome P450s (Supplementary Table 6). This pattern is consistent with an environmentally induced detoxification response in the lowland environment, paralleling inducible P450-mediated xenobiotic metabolism described for nectar and honey phytochemicals (Johnson et al. 2012; Mao et al. 2013). In lowland-origin colonies moved upslope (LL vs LH, baseline LL), we detected 50 DEGs (21 higher in LL; 29 higher in LH; median |log2FC| 1.37). The LH-upregulated set showed strong enrichment for chemosensory functions, particularly odorant binding, reflecting sensory retuning at higher altitude. This is congruent with the expanded odorant-receptor repertoire and chemosensory specialization of honey bees (Robertson and Wanner 2006). The native comparison between highland and lowland colonies at their home sites (HH vs LL, baseline LL) revealed 57 DEGs (20 higher in LL; 37 higher in HH; median |log2FC| 1.38). Here, genes related to cuticle and tegument were prominent: Apidermin-1 was among the most strongly up-regulated transcripts in HH, and multiple CPR (cuticular proteins) genes also differ between lineages, consistent with divergence in cuticle composition, sclerotization, and possibly thermal and desiccation properties.

Intersection analyses (UpSet; axis coding: 1 = HH vs HL, 2 = HH vs LL, 3 = LL vs LH) showed that these responses were largely contrast-specific, with limited sharing of individual DEGs. Considering all DEGs (up-and down-regulated combined), single-contrast sets dominated (1:42; 3:39; 2:35), whereas pairwise overlaps were modest (1-2:13; 2-3.8; 1-3:2), and only one gene was shared across all three contrasts. Directional analyses further clarified this structure: among up-regulated genes, single-contrast contributions prevailed (1:36; 3:13; 2:12), with one notable pairwise intersection (2-3:8) linking the native HH vs LL difference to the LL vs LH transplant. This shared set was enriched for proteostasis/unfolded protein response (UPR) chaperone module that remains elevated when lowland colonies are moved upslope. Among down-regulated genes, uniqueness was even more pronounced (2:36; 3:28; 1:22) with only a single shared intersection (2-3:1), indicating that repressed modules during acclimation are highly lineage- and context-specific.

Plastic responses were therefore largely origin-specific: only three DEGs were shared between the two translocation contrasts (HH vs HL and LL vs LH), and these changed in opposite directions. However, where translocation DEGs intersected the native HH vs LL difference, expression consistently shifted toward the local profile: expression flipped toward the high state in HH > HL and maintained the lowland-biased state in LL > LH, indicating convergence on the destination habitat. Gene-level patterns aligned with these broad axes. Multiple P450s were among the strongest signals in HH > HL; *abaecin* was elevated in HL, suggesting co-induction of immune defenses, (CASTEELS et al. 1990); odorant receptors (e.g., Or9, Or2, Or107) featured prominently in LL > LH; and *Apidermin-1* and CPRs characterized in HH > LL. Pigmentation-linked modules also emerged within these axes: yellow and yellow-like genes, together with consistent cuticle genes (*Apidermin,* CPRs) in the native HH vs LL contrast, align with canonical tanning biochemistry, in which Laccase2- mediated phenoloxidase activity and yellow family genes are required for cuticle pigmentation and sclerotization (Wittkopp et al. 2002; Arakane et al. 2005; True et al. 2005). Collectively, these results indicate that highland-to-lowland translocation is dominated by detoxification and immune challenges, lowland-to-highland translocation elicits chemosensory retuning, and proteostasis/UPR and cuticle biology form shared, cross-context components of acclimation.

### Cross-referencing reciprocal translocation and *AmEbony* CRISPR findings

Head RNA-seq from the *AmEbony* knockout experiment provided a complementary, tissue-specific perspective that nonetheless recapitulated several themes observed in the translocation data. The *AmEbony* knockouts head DEGs were dominated by odorant-binding proteins (OBP1, OBP1, OBP2, OBP5, OBP6, all up-regulated) and were enriched for odorant binding and extracellular region, indicating a head-centric chemosensory and secreted-protein program.

As expected, given differences in tissue (head vs whole body), subspecies (*A. m. carnica* vs East African lineages), and perturbation type (targeted gene knockout vs environmental translocation), gene-by-gene overlap between the set of bees obtained with the CRISPR/Cas9 method and transplant DEGs was negligible. However, in line with the intersection results (Figure 3), both datasets prominently feature members of the same functional families and modules rather than identical transcripts, namely: OBP and odorant receptors (OR, chemosensory), heat shock proteins (HSP) and chaperones (cellular homeostasis and stress) and cuticle-related genes (Apidermin, CPR). Core enzymes of the dopamine branch of the melanin pathway (e.g. ple/*TH*, *Ddc*, *AmEbony*, *tan*) did not emerge among whole-body transplant DEGs, whereas the presence of yellow and yellow-like genes, together with a strong cuticle signature in the HH vs LL native contrast, is consistent with tanning-associated cuticle chemistry and mechanistically align with the dopamine-NBAD branch governed by *ebony/tan* (True et al. 2005; Massey and Wittkopp 2016). This provides a functional bridge linking pigmentation, neuromodulation, and cuticular properties. In parallel, the detoxification and immune emphasis in HH > HL closely mirrors the inducible P450 and immune programs triggered by phytochemicals in honey and nectar (Johnson et al. 2012; Mao et al. 2013), reinforcing a view in which altitude-associated acclimation and pigment-pathway variation converge on chemosensation, cellular homeostasis and stress response (HSPs. UPR), detoxification/immune function (P450s, antimicrobial peptides), and cuticle biology (Apidermin/CPRs) rather than on one-to-one correspondence in specific transcripts (Robertson and Wanner 2006; McMenamin et al. 2020).

**Figure 3.**
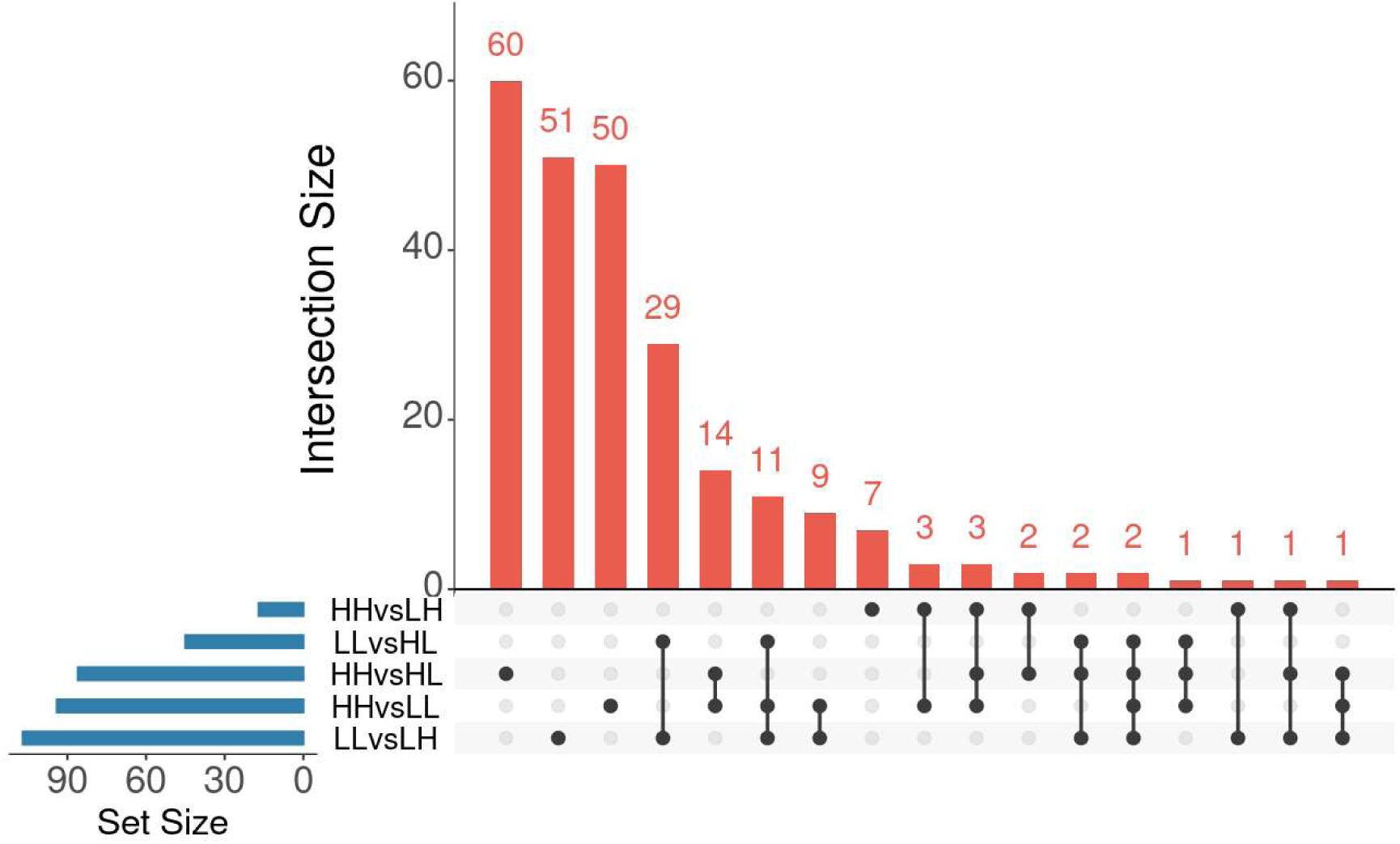
Overlap of candidate genes across translocation contrasts (UpSet plot). The UpSet plot summarizes genes identified in each pairwise comparison (HHvsLH, LLvsHL, HHvsHL, HHvsLL, and LLvsLH). Horizontal bars (blue) indicate the total number of genes per comparison (set size), while vertical bars (red) report the number of genes in each intersection. The dot matrix denotes which comparisons contribute to a given intersection (filled dots connected by lines), highlighting genes shared among multiple contrasts as well as genes unique to individual comparisons. Numbers above bars indicate intersection sizes.

## Discussion

East African honey bees occupy steep elevational gradients in which temperature, humidity, and solar irradiance change predictably over short spatial scales. Such systems provide a powerful natural laboratory for testing how pigmentation variation aligns with thermal ecology and how genome structure constrains or facilitates adaptation. Integration of population sampling across multiple mountain systems was employed to conduct a GWAS of abdominal pigmentation. This approach also involved structural-variant genotyping of two large altitude-associated haplotype blocks (r7, r9) and CRISPR/Cas9-induced disruption of *ebony* gene function followed by transcriptome- and network-level analyses. By this strategy we aimed to establish a connection between clinal phenotype, genomic architecture, and causal perturbation. The integrated design enabled us to investigate the following research questions: first, whether a limited set of loci can exert disproportionately large phenotypic effects against a polygenic background, and second, whether pleiotropy links pigmentation to chemosensory, metabolic, and redox biology (Clusella - Trullas et al. 2008; Boyle et al. 2017; Clusella-Trullas and Nielsen 2020).

### A top-heavy polygenic architecture centered on *AmEbony*

Our GWAS revealed a broadly polygenic signal for abdominal pigmentation, but with a striking concentration of association around a single locus: *AmEbony*. This pattern is characteristic of a “top-heavy polygenic” architecture, in which a small number of variants explain a large fraction of the variance while many additional loci make individually minor contributions. Such architectures reconcile the presence of strong, interpretable association peaks with the long polygenic tails expected for quantitative traits and are consistent with core-plus-peripheral or omnigenic models, in which peripheral networks modulate the effects of core pathway genes (Boyle et al. 2017). Population genomic surveys in *A. mellifera* have documented extensive standing genetic variation and repeated signatures of local adaptation along environmental gradients (Gruber et al. 2013; Wallberg et al. 2017). Against this background, our finding that pigmentation variation is overwhelmingly concentrated at *AmEbony*, with additional sub-genome-wide peaks, fits a scenario in which selection operates primarily on key nodes in the melanin pathway while also fine-tuning trait expression via a broader network of small-effect loci.

### A small fraction of SNPs explains disproportionate phenotypic variance

The observation that a modest set of lead SNPs at *AmEbony* captures most of the variance in abdominal pigmentation is not paradoxical. Melanin synthesis is organized around a few biochemically central reactions, including the conjugation of dopamine with β-alanine by *ebony*. Variants that modify expression or function of such core enzymes (or their immediate regulators) can have large optical and thermal consequences even when the underlying allelic changes are subtle. Our data show exactly this pattern: a dense block of highly associated variants at *AmEbony*, including missense, UTR, and intronic SNPs, segregates in three distinct genetic clusters that map almost perfectly onto the phenotypic pigmentation gradient. At the same time, the polygenic background remains detectable, in line with core-plus-peripheral models of complex traits (Boyle et al. 2017).

Importantly, the direction of effect is consistent with the Thermal Melanism Hypothesis: darker alleles at *AmEbony* are enriched at higher elevations, implying thermoregulatory benefits under cooler highland conditions, consistent with macro-physiological tests across insects (Clusella Trullas et al. 2007; Boyle et al. 2017; Clusella-Trullas and Nielsen 2020). Thus, *AmEbony* functions as a major-effect locus that channels substantial adaptive response along the elevational gradient, embedded within a polygenic architecture.

### r7 and r9 inversion polymorphisms are associated with pigmentation, but likely act via genomic background

We confirm that the large haplotype blocks r7 and r9, previously inferred as inversions and tied to high-altitude adaptation (Wallberg et al. 2017; Mazzoni et al. 2025), are present not only in previously studied regions but also on Mount Kilimanjaro and Mount Meru. This extends the geographic range of these structural variants and strengthens the case that they are repeatedly recruited in high-elevation ecotypes. Inversion genotypes at r7 and r9 are non-randomly associated with pigmentation clusters: inverted (highland) haplotypes are overrepresented among darker bees, whereas the r5 region shows no such pattern. This suggests that r7 and r9 do not directly encode pigmentation changes but rather maintain highland genomic backgrounds on which dark *AmEbony* alleles segregate. Consistent with theory (Kirkpatrick and Barton 2006), these inversions likely preserve co-adapted multi-gene complexes that include, but are not limited to, pigmentation. *AmEbony* thus provides a mechanistic link to melanism, while r7 and r9 serve as structural scaffolds for broader highland adaptation.

### CRISPR/Cas9 establishes *AmEbony* as a causal pigmentation locus in *Apis mellifera*

Our CRISPR/Cas9 experiments in *A. m. carnica* provide direct functional evidence that *AmEbony* plays a causal role in honey bee pigmentation. Targeted disruption of *AmEbony* produced clear pigmentation changes, consistent with the locus acting as a key node in melanin biosynthesis and strongly supporting its identification as the major GWAS peak. Although rescue experiments are not possible in the honey bee yet, our results align with prior CRISPR-based manipulations of pigmentation and sensory genes in *A. mellifera* (e.g. *yellow* and *orco*). The feasibility of genome editing in *A. mellifera* moves the field beyond correlative association toward mechanistic dissection, enabling tests of pleiotropy versus linkage and functional validation of candidate loci underpinning ecological adaptation (Hu et al. 2019; Chen et al. 2021; Nie et al. 2021).

### Gene expression and network analyses link pigmentation, chemosensation, and redox physiology

Head transcriptomes from *AmEbony* knockouts showed consistent shifts in modules enriched for odorant binding and oxidoreductase activity. These results are mechanistically coherent: in *Drosophila*, *ebony* is expressed in glia and cuticle, controls histamine/dopamine conjugation, and couples tanning, neuromodulation, and behavior; *ebony/tan* edits reshape cuticular hydrocarbons, which act as chemosensory cues (Hovemann et al. 1998; Hartwig et al. 2014; Izoré et al. 2019; Massey et al. 2019). In honey bees, an expanded odorant-receptor repertoire and specialized receptors for queen pheromones highlight the centrality of olfaction (Robertson and Wanner 2006; Wanner et al. 2007; Vernier et al. 2019). We therefore propose a pleiotropy-first model: *Ebony*-mediated changes in dopamine/biogenic-amine metabolism and/or CHC composition alter neuromodulatory tone and chemical context at the body surface, which in turn influence odorant-receptor expression and olfactory circuit function. Our GENIE3-based network analyses support this view: oxidoreductase activity (GO:0016491) was enriched in 5/5 independent runs, and odorant binding (GO:0005549) in 4/5, indicating robust involvement of redox and chemosensory pathways, whereas most other GO terms were run-sensitive and are best treated as hypotheses. The redox signal aligns with the oxidative chemistry of melanization and cuticle tanning and with known links between mitochondrial function, thermal performance, and energy metabolism in bees.

### Reciprocal translocation highlights convergent physiological axes of acclimation

Reciprocal translocations revealed that highland-to-lowland and lowland-to-highland moves elicit distinct but convergent physiological responses. Highland-origin colonies moved to lowland showed strong induction of cytochrome P450s and monooxygenase activity, consistent with environmentally triggered detoxification and immune programs in response to novel phytochemicals and xenobiotics (Johnson et al. 2012; Mao et al. 2013). Lowland-origin colonies moved upslope upregulated odorant receptors and odorant-binding proteins, suggesting chemosensory retuning to high-altitude conditions. This is precisely the functional axis which was also highlighted in *AmEbony* head knockouts (Robertson and Wanner 2006).

The native HH vs LL comparison emphasized cuticle biology (Apidermin-1, CPR cuticular proteins) and pigmentation-linked genes (*yellow*, *yellow*-*like*), consistent with integrated evolution of cuticle structure, coloration, and function (Wittkopp et al. 2002; Arakane et al. 2005; True et al. 2005). Intersection analyses showed minimal sharing of individual DEGs across contrasts, but strong convergence at the level of functional modules: chemosensation (OBPs/ORs), detox/immune (P450s, antimicrobial peptides such as *abaecin*) (CASTEELS et al. 1990), proteostasis and stress (heat-shock/chaperone pathways) (McMenamin et al. 2020),and cuticle/Apidermin biology (including emerging antimicrobial roles); (Kim et al. 2022). Expression shifts in transplanted colonies tended to move toward the local native profile, indicating convergence on destination-specific expression states. *AmEbony* and the conserved dopamine-NBAD branch of melanization (Wittkopp et al. 2002; Wittkopp et al. 2003; True et al. 2005) provide a mechanistic nexus linking pigmentation chemistry to neuromodulation, cuticle properties, and the same chemosensory and homeostatic axes that are environmentally modulated by elevation.

In summary, our study shows how a canonical pigmentation gene (*AmEbony*), embedded in a top-heavy polygenic architecture and interacting with large inversion polymorphisms and environmentally responsive regulatory networks, underlies adaptive thermal melanism in a eusocial pollinator.

## Materials and Methods

### Whole Genome sequencing and analysis

DNA was extracted from 72 full thorax from one honey bee per colony using the CTAB protocol (Rusterholz et al. 2015) and processed to 2 x 150bp paired-end sequencing (Illumina Novaseq X) targeting 25x coverage (Biomarker Technologies (BMK) GmbH). To get a more comprehensive Genome Wide Association Study (GWAS) concerning the pigmentation of honey bees we integrated the newly sequenced samples in a larger dataset including additional East African honey bee publicly available whole genome data (Short Read Archive (SRA). We included 24 highland and lowland honey bees collected in Uganda sequenced with a mean coverage of 25x (Mazzoni et al. 2025) (Bioproject: PRJNA1179698), and 39 highland and lowland honey bees sampled in Kenya in two different regions (Wallberg et al. 2017) (Bioproject: PRJNA357367), all with 12x mean coverage. SNP calling was carried with the very same pipeline as in (Mazzoni et al. 2025). We used ‘HaplotypeCaller’ from gatk4 v4.3.0.0, followed by joint genotyping using the ‘GenotypeGVCFs’ function (der Auwera et al. 2013; Poplin et al. 2017). We imported data in a batch size of 20 by chromosome with ‘GenomicsDBImport’. The resulting VCFs were merged with bcftools v1.11 (Danecek et al. 2021). Allele balance for each variant was annotated with gatk3 v3.8.1.0 and filtered using vcffilter with the condition ‘ABHet > 0.25 & ABHet < 0.75 | ABHet < 0.01‘, to allow for homozygous variants, too (Garrison et al. 2022). After evaluating the distribution of the parameters from the INFO field in the final VCF, additional filters were applied with bcftools view v1.11 (Danecek et al. 2021). The filtering criteria for variant sites included the following thresholds: MQ < 40, MQRankSum < -5, MQRankSum > 5, ExcessHet > 20, AC = 1, INFO/DP < 11660, INFO/DP > 21654, AN < 826, FS > 20, SOR > 5, and QD < 5. Only biallelic SNPs were considered, and indels were excluded. The gatk4 function ‘SelectVariants’ was used to retain only variant sites in chromosomes for downstream analysis (der Auwera et al. 2013). Additionally, samples with a coverage of less than 6x were removed from the dataset. Finally, the remaining variants were phased using beagle v5.4 (22Jul22.46e.jar) (Browning et al. 2018; Browning et al. 2021).

### Genome Wide Association study and population genomics statistics

A genome-wide association study (GWAS) was performed to investigate the genetic basis of variation in abdominal pigmentation. The analysis used both the whole-genome sequencing (WGS) data generated in this study and publicly available genomic data for the same species. GWAS was carried out using GEMMA (Zhou and Stephens 2012) (Genome-wide Efficient Mixed Model Association), which implements a univariate linear mixed model (LMM) framework. This approach incorporates a genetic relatedness matrix (GRM) as a random effect to account for population structure and cryptic relatedness, thereby reducing confounding and improving the reliability of the association signals. Each single-nucleotide polymorphism (SNP) was tested individually for its association with the pigmentation phenotype using the LMM as implemented in GEMMA. Statistical significance was evaluated using the Wald test, and the resulting p-values were adjusted for multiple testing using the Benjamini–Hochberg false discovery rate (FDR) procedure, with a significance threshold of FDR < 0.05. To further characterize genomic variation in relation to pigmentation phenotypes, we computed standard population genetic statistics nucleotide diversity (π), fixation index (F_ST_), and absolute genetic divergence (D_XY_) using pixy (Korunes and Samuk 2021). Calculations were performed with the flags --stats pi fst dxy, assuming a window size of 1 bp, thereby enabling site-wise resolution.

### CRISPR/Cas9 experiment design

The phylogenetic tree to assess the *AmEbony* orthology to the *D. melanogaster* ebony gene was performed using iQtree2 running on 12 threads performing ultrafast bootstrap with 1000 replicates and allowing the tool to find the best evolutionary model with the -m MFP flag (Minh et al. 2020). The honey bee pigmentation-related gene *ebony* (NCBI: LOC409109/GB46429) is located on chromosome 1 and comprises 15 exons. We compared the three NCBI transcript variants and found that they encode identical amino acid sequences. Therefore, we used the longest transcript (XM_026445876.1) for sgRNA target selection. Candidate crRNA targets were identified in Benchling (Benchling 2023) (Benchling 2023) following (Değirmenci et al. 2020): 20-bp protospacers with an NGG PAM and a 5′ guanine. We prioritized guides with high predicted cutting efficiency (on-target >50) and high specificity (low off-target risk). Candidate sites were additionally validated with SPLIGN (Kapustin et al. 2008) to ensure they lay entirely within an exon and with Vienna RNAfold (Lorenz et al. 2011) to confirm appropriate sgRNA secondary structure and accessibility for Cas9 interaction. To maximize functional disruption, we favored targets positioned early in the *ebony* open reading frame (within ∼800 bp of a 2583-bp ORF). The three best guides were synthesized in-house via overlapping Phusion PCR to generate the DNA template, followed by T7 in vitro transcription, sgRNA purification, and quantification as described in (Değirmenci et al. 2020). Pilot injections across multiple sgRNA concentrations were used to estimate both larval hatching success and mutation frequency, which were combined into a “mutant rate” to rank treatments. The best-performing guide (5′-GCAGGAATTACCTTATGCAG-3′; ORF position 635 on the plus strand) was selected for the main experiment at 92 ng/µl sgRNA with 3.13 µM Cas9, delivering 400 pl per embryo. Embryos were collected from *Apis mellifera carnica* colonies using Jenter systems and injected within 0-1.5 hours after oviposition to target the single-cell stage. Artificial rearing of the honey bees in the laboratory followed established protocols (Schmehl et al. 2016; Değirmenci et al. 2020). Adult workers were phenotyped at five days post-emergence and mutations were confirmed by fluorescently labeled PCR spanning the target site and capillary electrophoresis, comparing amplicon length shifts to the 171-bp wild-type fragment.

### Reciprocal translocation experiment

To investigate the transcriptional responses of honey bees to environmental change across elevations, we conducted a reciprocal translocation experiment. A total of four colonies originating from lowland labelled LL (for lowland controls) and LH (for lowland translocated to highland colonies) sites were relocated to highland locations, while five colonies from highland sites HH (for highland controls) and HL (for highland translocated to lowland) were moved to lowland environments. At each location, we also maintained a set of control colonies that were kept in their original environment and sampled in parallel with the translocated colonies. These control hives provided an essential baseline for disentangling the effects of translocation from local environmental variation. The experiment was carried out in collaboration with local farmers, who hosted the colonies at the designated sites. Regular monitoring and colony maintenance were ensured by our partners at the University of Embu, who performed scheduled inspections throughout the course of the study. To preserve colony identity and prevent potential queen dispersal during the experiment, queen-excluder grids were installed at the entrances of all translocated hives. The experiment was conducted over a period of 9 months from June 2023 to February 2024, allowing us to capture the longer-term transcriptional responses of honey bees to sustained changes in environmental conditions at both high and low elevations. At the end of the period, RNA was extracted from heads of 3 honey bees per beehive and sequenced (see below).

### RNA extraction library preparation and sequencing

For the functional transcriptomic analysis of *AmEbony*, we collected 12 individual bees, comprising 6 Eby⁺/Eby⁺ (wild-type) and 6 Eby⁻/Eby⁻ (mutant) individuals. From each bee, we dissected and processed three distinct tissues: head, thorax, and abdomen. To investigate potential tissue-specific transcriptional responses to the loss of *AmEbony* function. Total RNA was extracted from each tissue sample using TRIzol reagent (Thermo Fisher Scientific) according to the manufacturer’s guidelines with minor optimizations for honey bee tissues. In brief, each sample was homogenized in 500 μL of TRIzol, followed by phase separation with 100 μL of chloroform and precipitation with 300 μL of isopropanol. After centrifugation and solvent removal, the RNA pellet was resuspended in 30 μL of RNase-free water and stored at −80 °C until further processing. The integrity and concentration of the RNA were assessed prior to library preparation followed by standard barcoding procedures for multiplexing and sequencing on an Illumina NovaSeq 6000 platform with a 2 × 150 bp paired-end configuration. Each library generated approximately 40 million high-quality reads per sample, ensuring sufficient coverage for quantitative transcriptome profiling across all three tissue types. The same extraction and sequencing procedure were applied to honey bees involved in the translocation experiment, but here we extracted and sequenced RNA only from head of the animals.

For transcriptome sequencing the reads quality was checked using fastqc (Andrews 2010). Adapters were removed and reads were trimmed using Trimmomatic v0.36 (Bolger et al. 2014). After trimming, fastqc was used again to check the quality of the trimming process. Trimmed reads were used by Kallisto (Bray et al. 2016) to quantify the abundances of transcripts. Then, DESeq2 (Love et al. 2014) was used to analyse differentially expressed genes. Finally, a GO enrichment analysis was done with gProfiler2 (Kolberg et al. 2020; Kolberg et al. 2023).

### Gene regulatory network construction and analysis (GENIE3)

To investigate the transcriptional architecture associated with AmEbony function, we constructed a gene regulatory network (GRN) using the GENIE3 algorithm (Huynh-Thu et al. 2010). The network inference was performed on the variance-stabilized expression matrix derived from the RNA-seq data of Eby⁺/Eby⁺ (wild-type) and Eby⁻/Eby⁻ (mutant) individuals. Prior to network construction, genes with consistently low expression were removed, and the remaining genes were variance-stabilized to reduce heteroscedasticity across samples. GENIE3 was executed using a fixed random seed to ensure reproducibility, with 2,000 trees per random forest (nTrees = 2000) and leveraging 12 computational cores (nCores = 12) to optimize performance. The algorithm produced a weighted adjacency matrix, where each edge weight reflects the predicted importance of a potential regulator gene for the expression of a target gene. To focus on the most robust interactions, only the top 0.05% of edges corresponding to those with weights above the 99.95th percentile of the weight distribution was retained. This high-confidence edge set was used to construct a directed graph representing the core transcriptional network inferred by GENIE3.

### Ebony gene tree

The ebony gene tree of Hymenoptera order was inferred using IQtree2 (Huynh-Thu et al. 2010) allowing the tool to find the best model (-MFP flag) and allowing for ultra-fastbootstrap (-bb flag) with 1000 replicates. The accessions used to build the tree are reported in Supplementary Table 7.

## Data availability

All newly sequenced honey bee genomes and transcriptomes were deposited on SRA with the bioproject accession number: PRJNA1445831.

## Acknowledgments

We would like to thank our local partners in Tanzania and Kenya, and the extraordinary work carried out by our field work assistants, Kennedy in Kenya, and Mark in Tanzania, as well as all the local communities we came in touch with during our trips to East Africa.

This research was supported by grants of the Deutsche Forschungsgemeinschaft to M.H. and R.S. (HA5499/11-1; SCHE 1573/14-1) and by a grant of the VW Foundation to R.S. (VW Momentum).

