## Supplementary Information for "A Polygenic Route to Thermal Melanism and high-elevation adaptation in Honey Bees"

**This PDF file includes:**

Legends for Supplementary Figure S1 to S8

Legends for Supplementary tables S1 to S7

Dataset link for SRA

figures

Fig. S1.

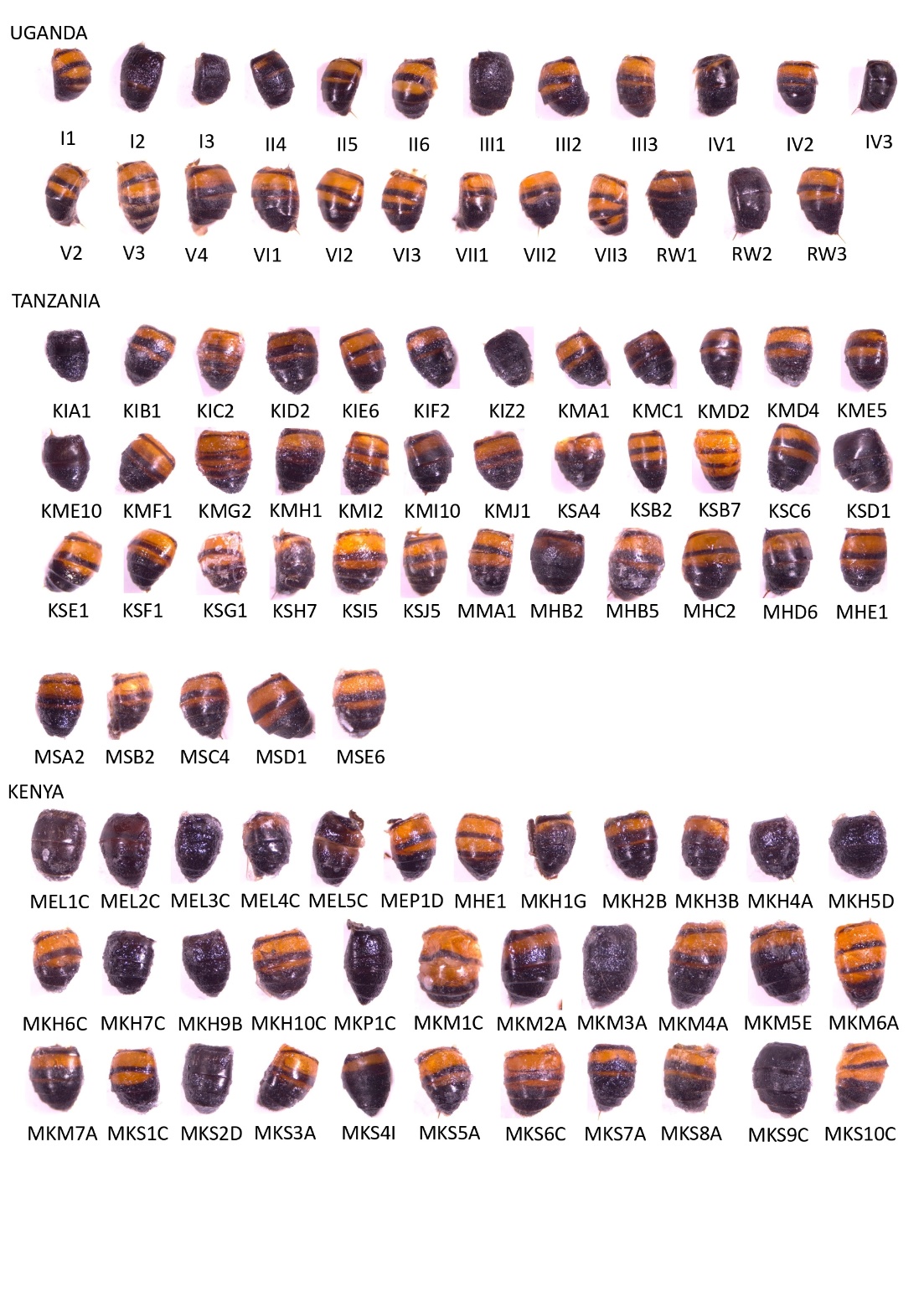

Photographs of the abdominal color patterns of all African honey bee specimens used in this study. The specimens shown here were sequenced and are organized by country and sample identifier.

Fig. S2.
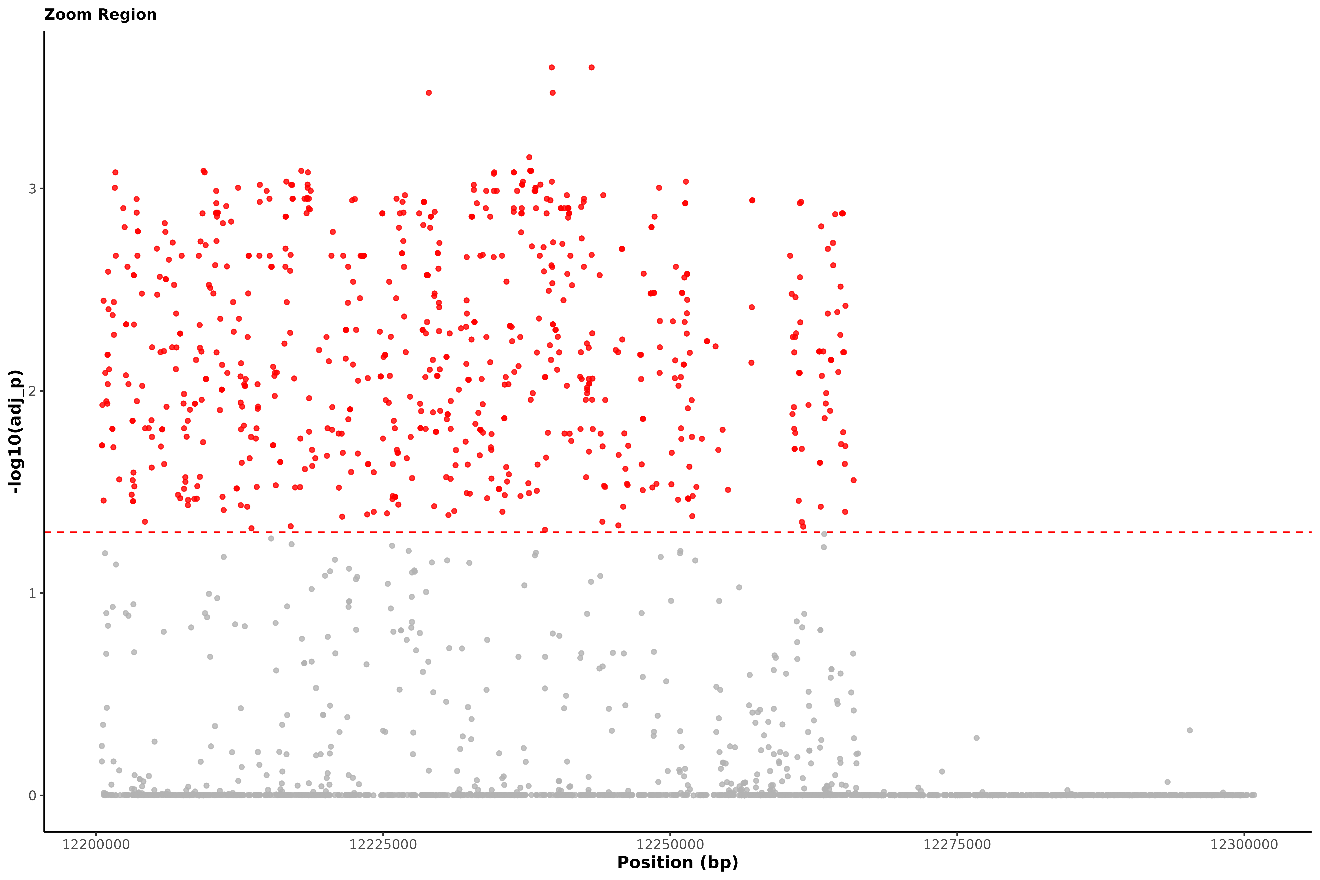
Zoomed-in view of the GWAS association signal in the genomic region containing the main peak of significance. Each point represents a SNP plotted by genomic position and log10 adjusted p-value; red dots indicate SNPs above the significance threshold.

**Fig. S3.**
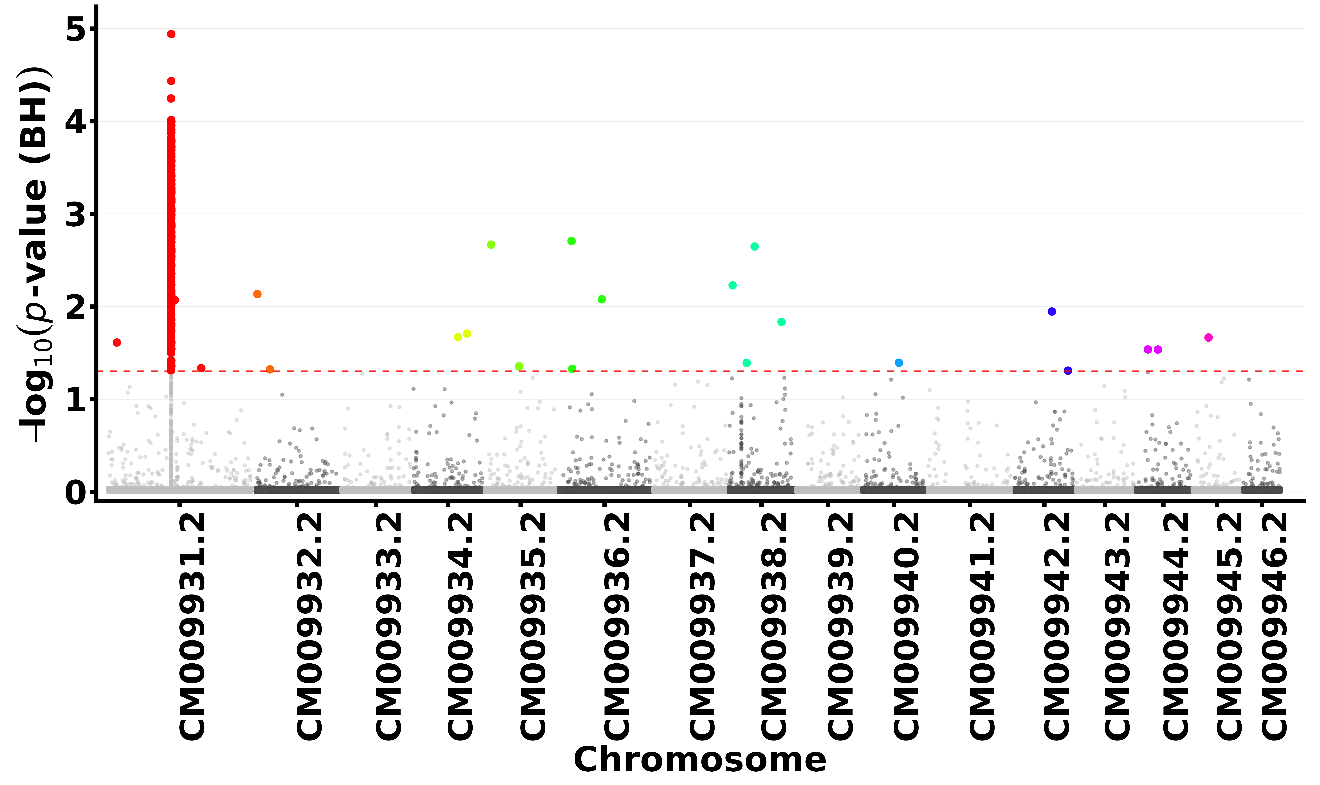
 Results of the leave-one-chromosome-out (LOCO) analysis used as an independent validation of the GWAS signal. The strongest associations remain concentrated in the same genomic region, supporting the robustness of the detected peak.

Fig. S4.
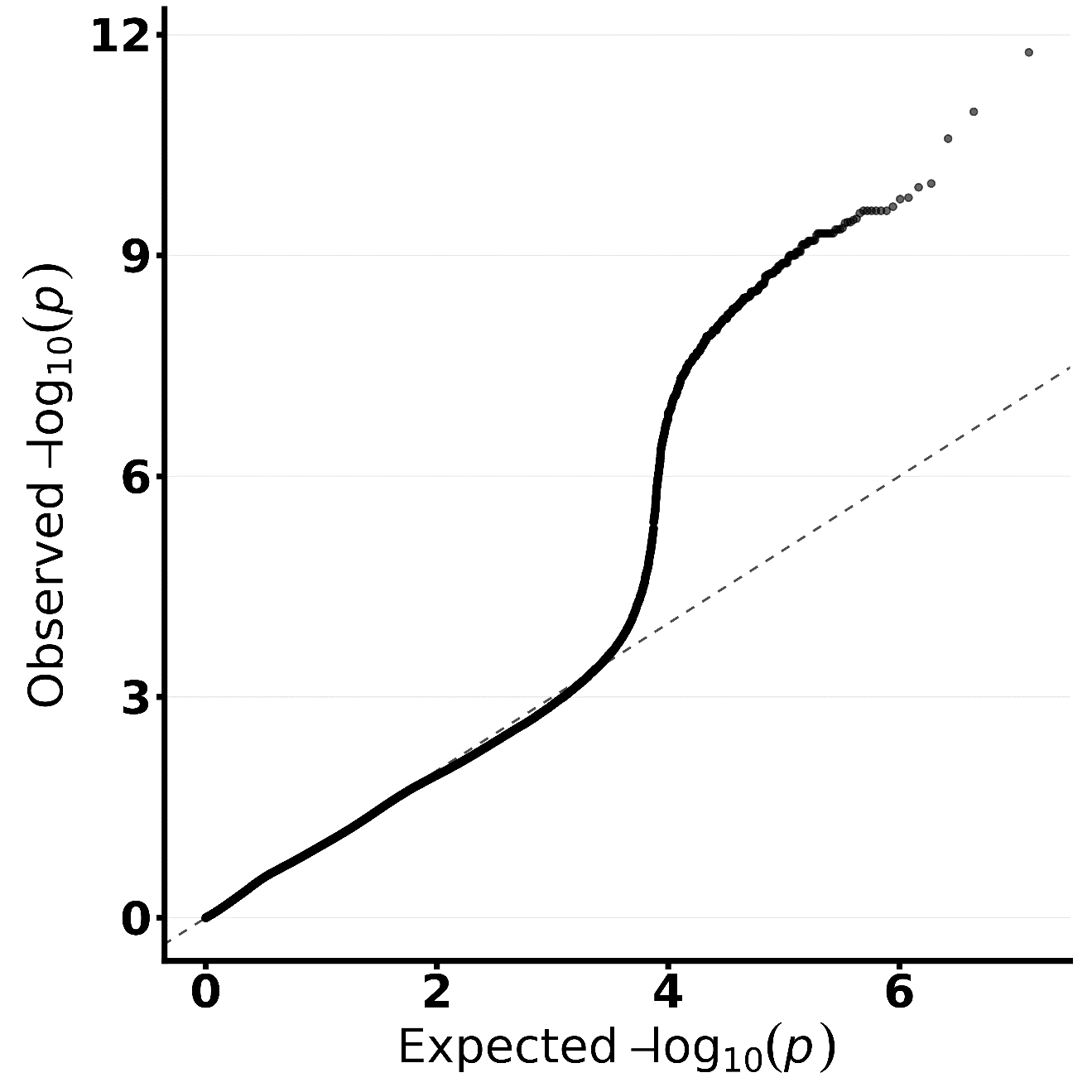
 QQ plot for the LOCO analysis used to assess the distribution of association test statistics. The departure of the observed values from the expected null distribution is consistent with the presence of true association signals.

Fig. S5.
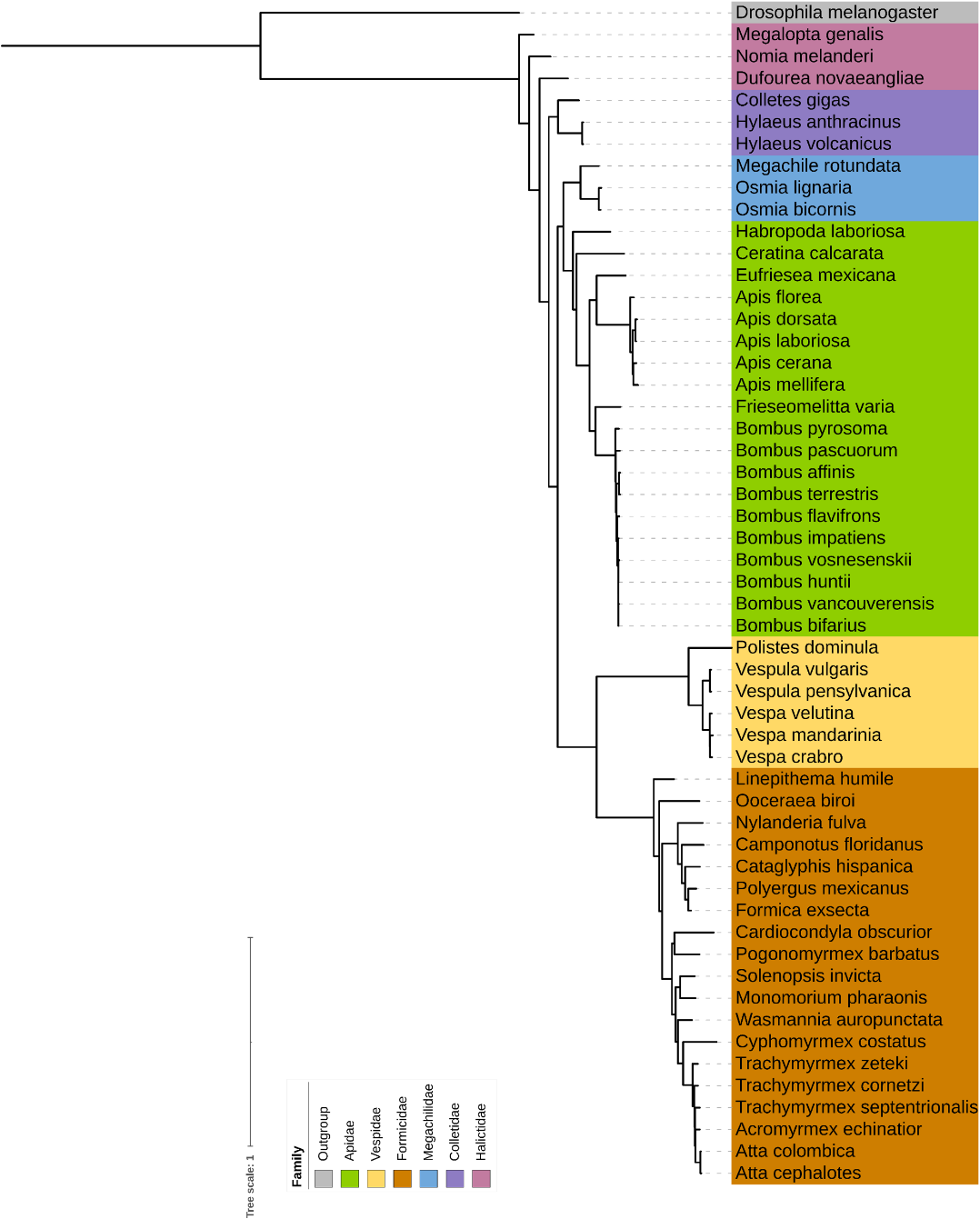
Gene tree of ebony orthologs across Hymenoptera, rooted with Drosophila melanogaster as the outgroup. Tip colors indicate clusters corresponding to different hymenopteran families.

Fig. S6.
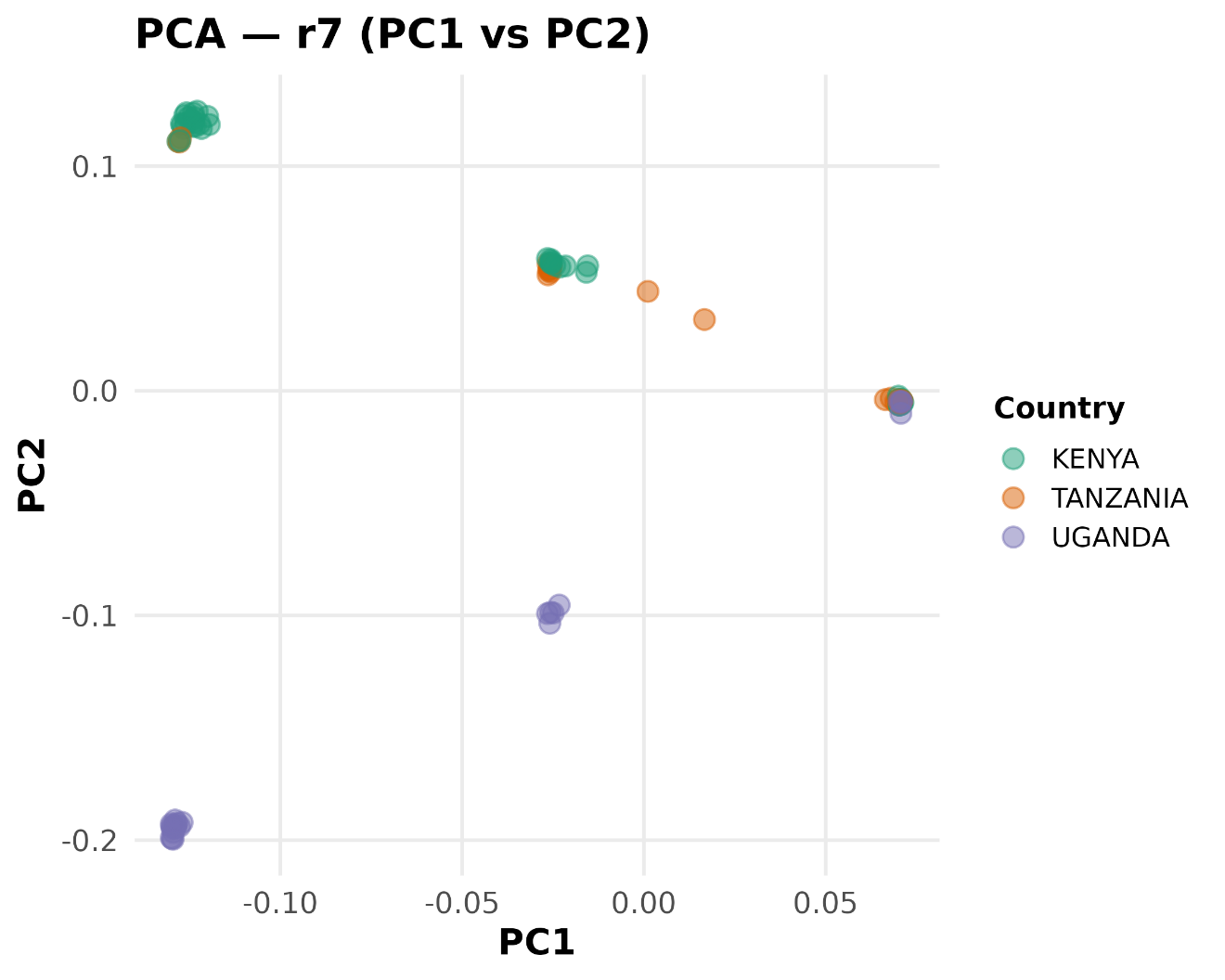
Principal component analysis of genetic variation across the r7 inversion region. Tanzanian samples cluster with inversion-associated groups, suggesting that this chromosomal inversion is also present in Tanzania.

Fig. S7.
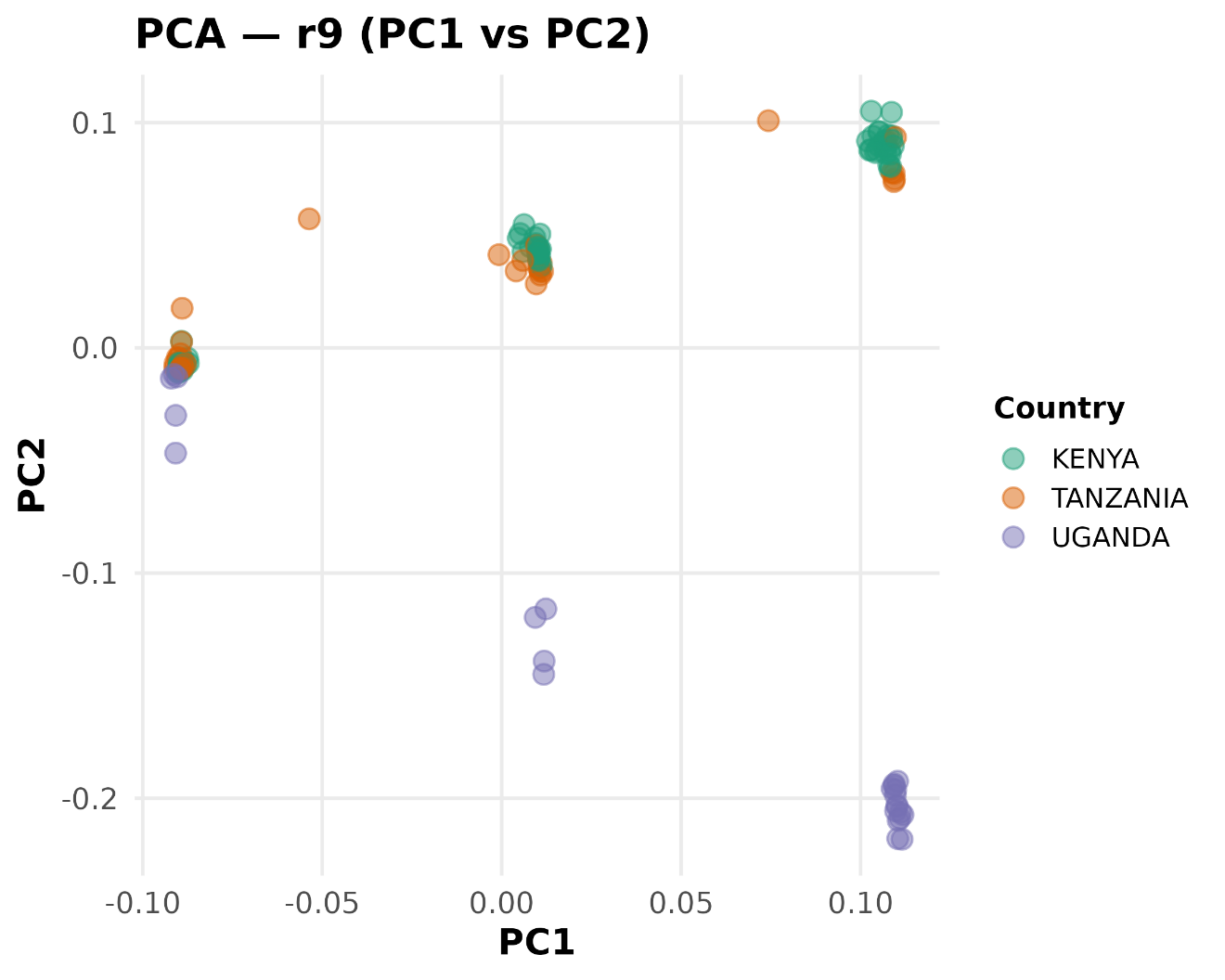
Principal component analysis of genetic variation across the r9 inversion region. Tanzanian samples cluster with inversion-associated groups, suggesting that this chromosomal inversion is also present in Tanzania.

Fig. S8..

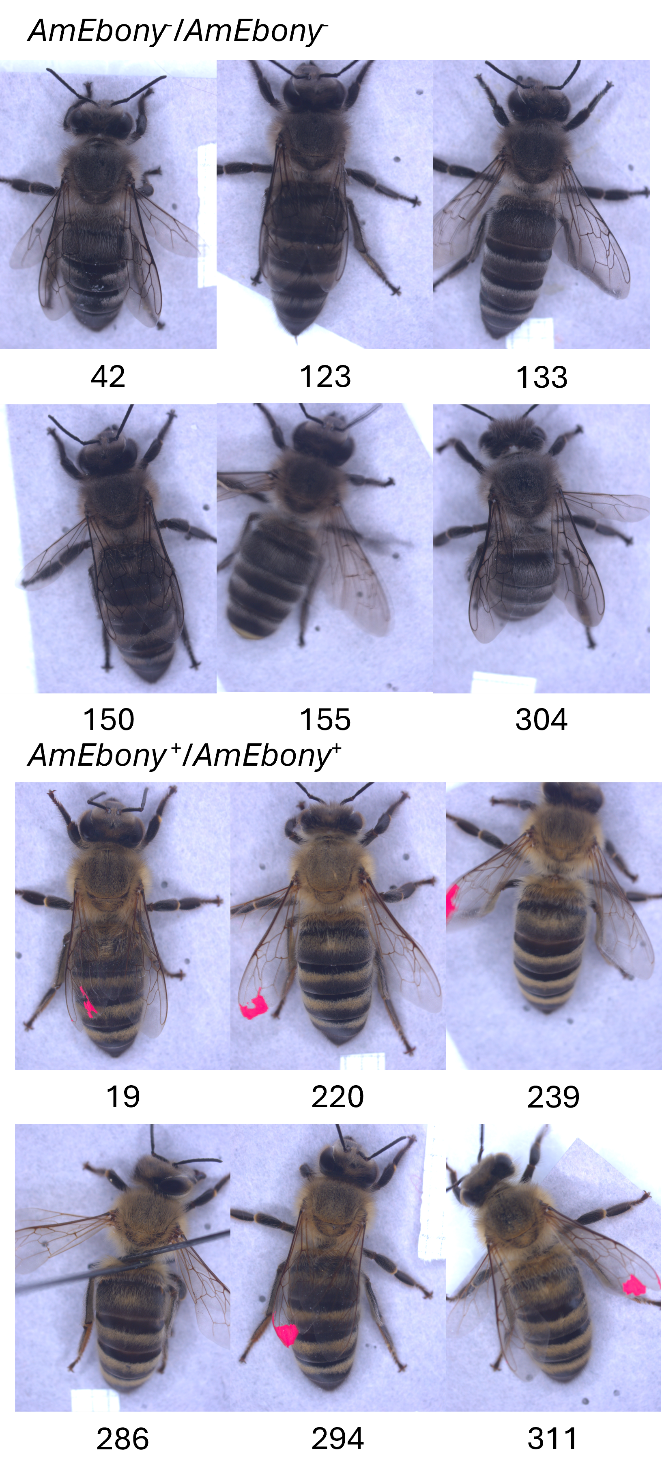

*Apis mellifera* individuals from the CRISPR/Cas9 experiment targeting *AmEbony* gene. The disruption of the gene resulted in markedly darker body pigmentation compared with wild type individuals

Tables

Supplementary Table 1. Coordinates of the sampled specimens and colonies from Kenya and Tanzania.

Supplementary Table 2. Pigmentation metadata of the dataset

Supplementary Table 3. Annotation of Single nucleotide polymorphisms (SNPs) associated with the dark pigmentation and annotated with SNPeff.

Supplementary Table 4. Association between inversions r7 and r9 and the pigmentation phenotype. The association was computed using Chi-square test.

Supplementary Table 5. Results of significant modules and relative genes of the different GENIE3 runs.

Supplementary Table 6. RNA-seq results for the comparison of translocated colonies with the relative controls, as well as the comparison between Highland and Lowland colonies.

Supplementary Table 7. Genes accession numbers and relative species and family information relative to the hymenopteran gene tree.

**SRA Dataset link (for reviewers)**

**https://dataview.ncbi.nlm.nih.gov/object/PRJNA1445831?reviewer=474ivo0uj4pmk5t49i4f2d893m**
