## Supplementary figures and images for "A Polygenic Route to Thermal Melanism and high-elevation adaptation in Honey Bees"

### Supplementary Figure 1

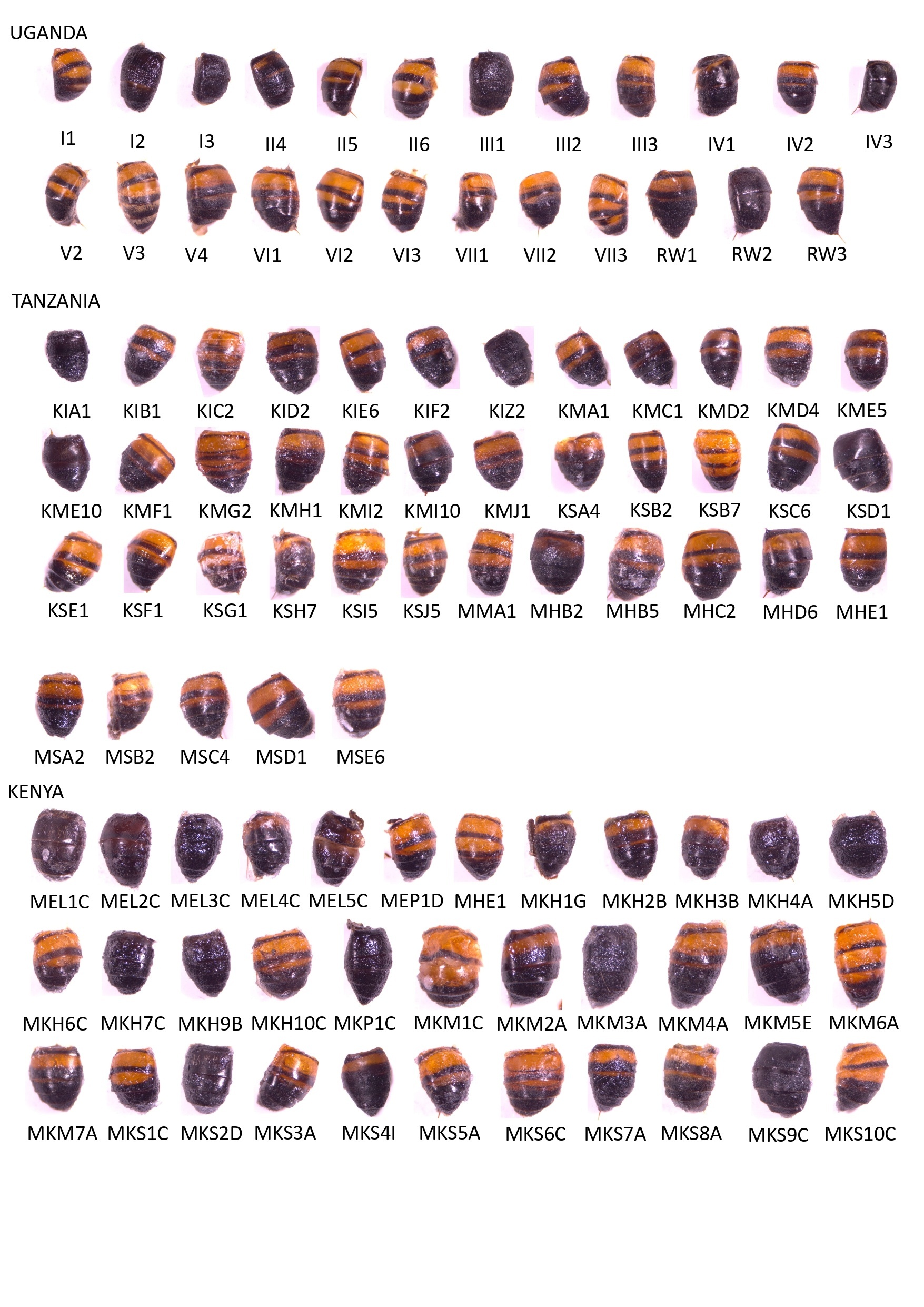

### Supplementary Figure 2

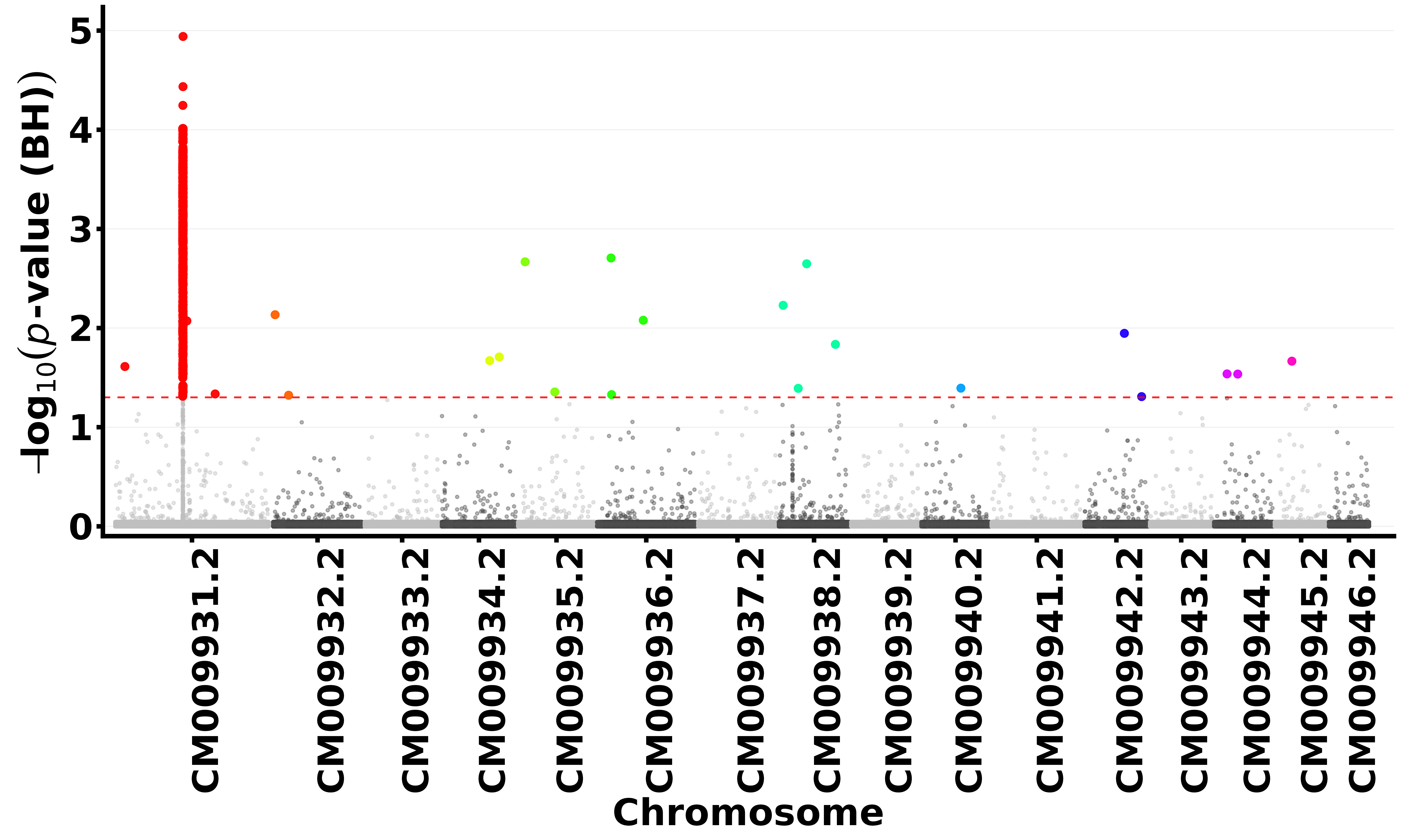

### Supplementary Figure 3

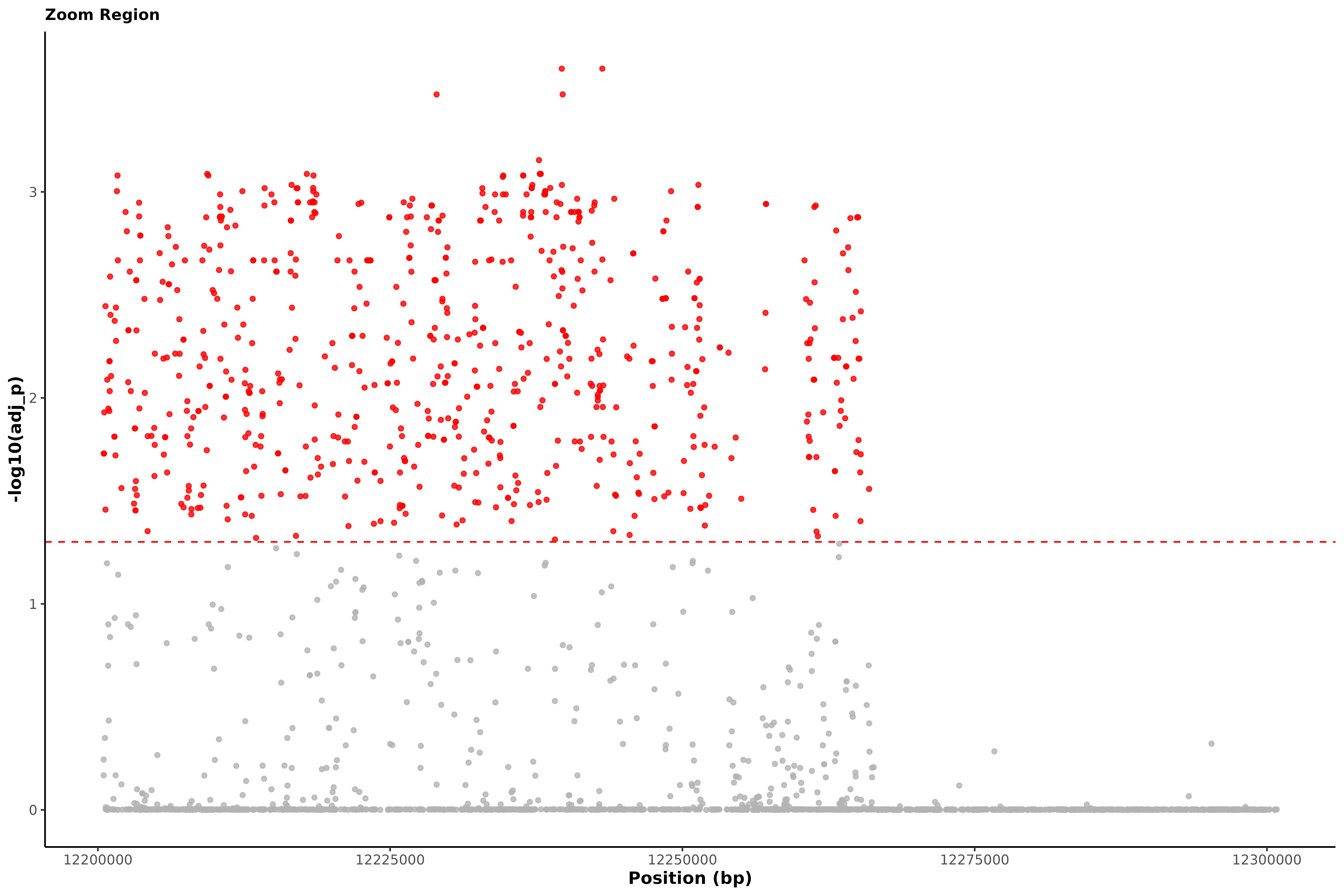

### Supplementary Figure 4

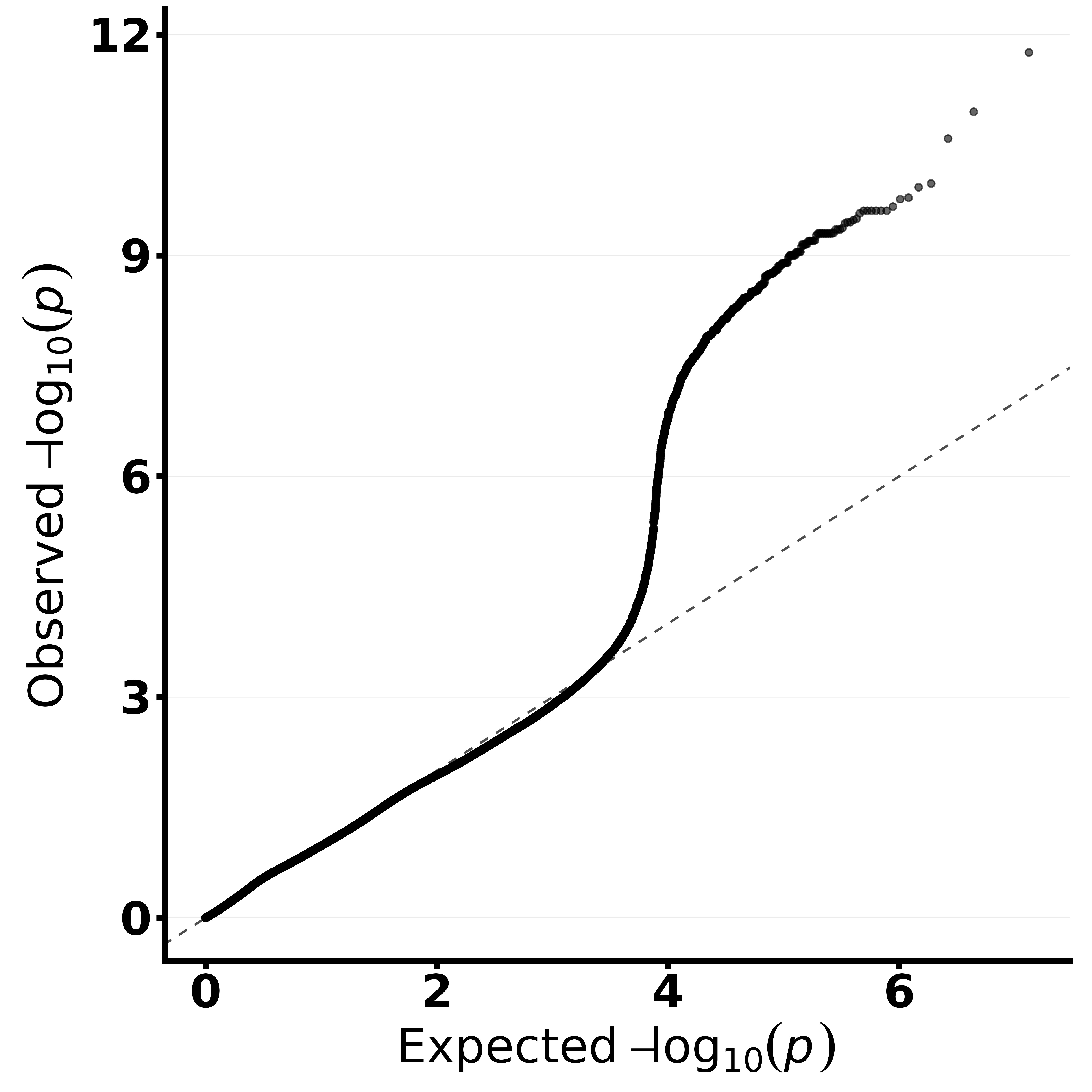

### Supplementary Figure 5

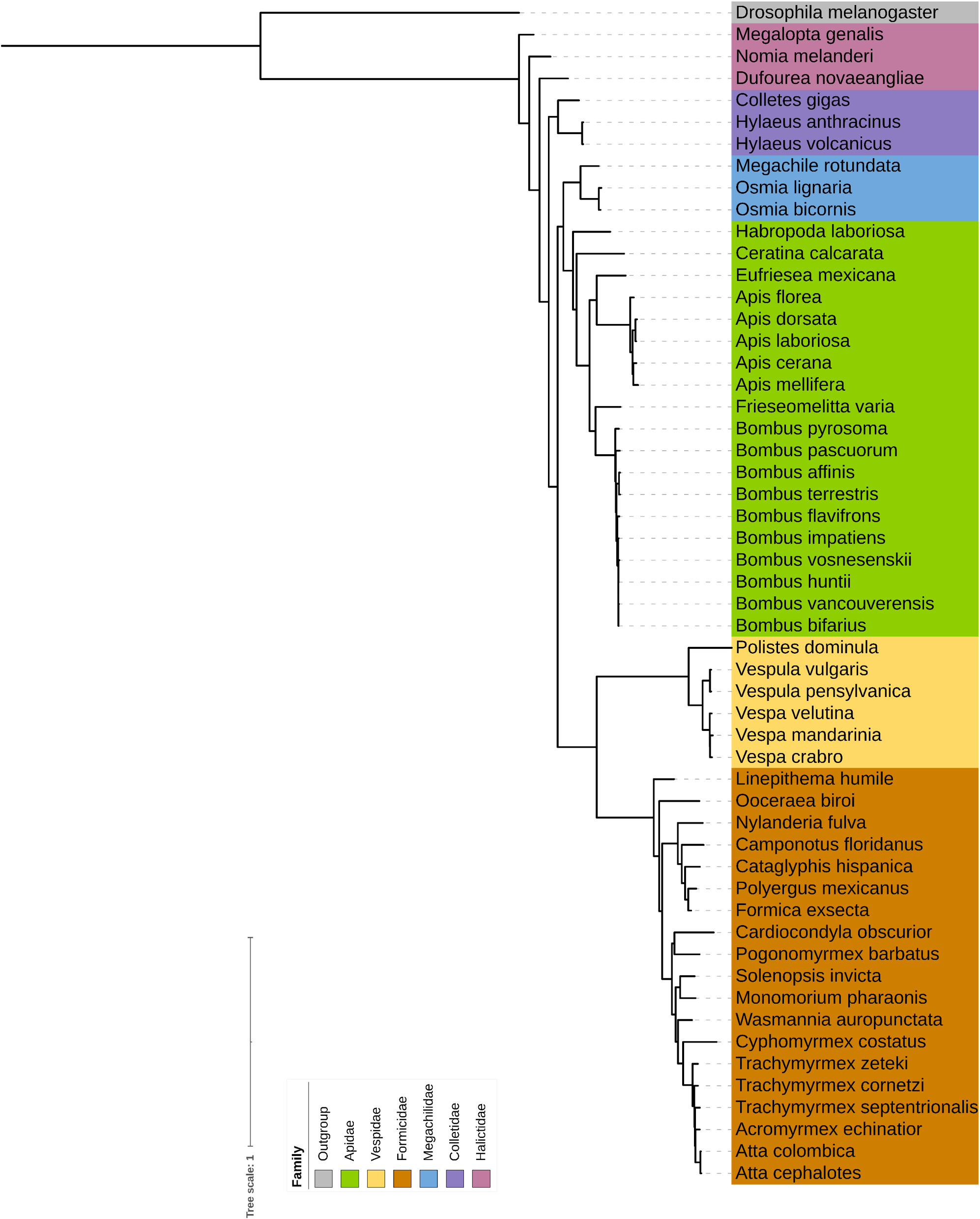

### Supplementary Figure 6

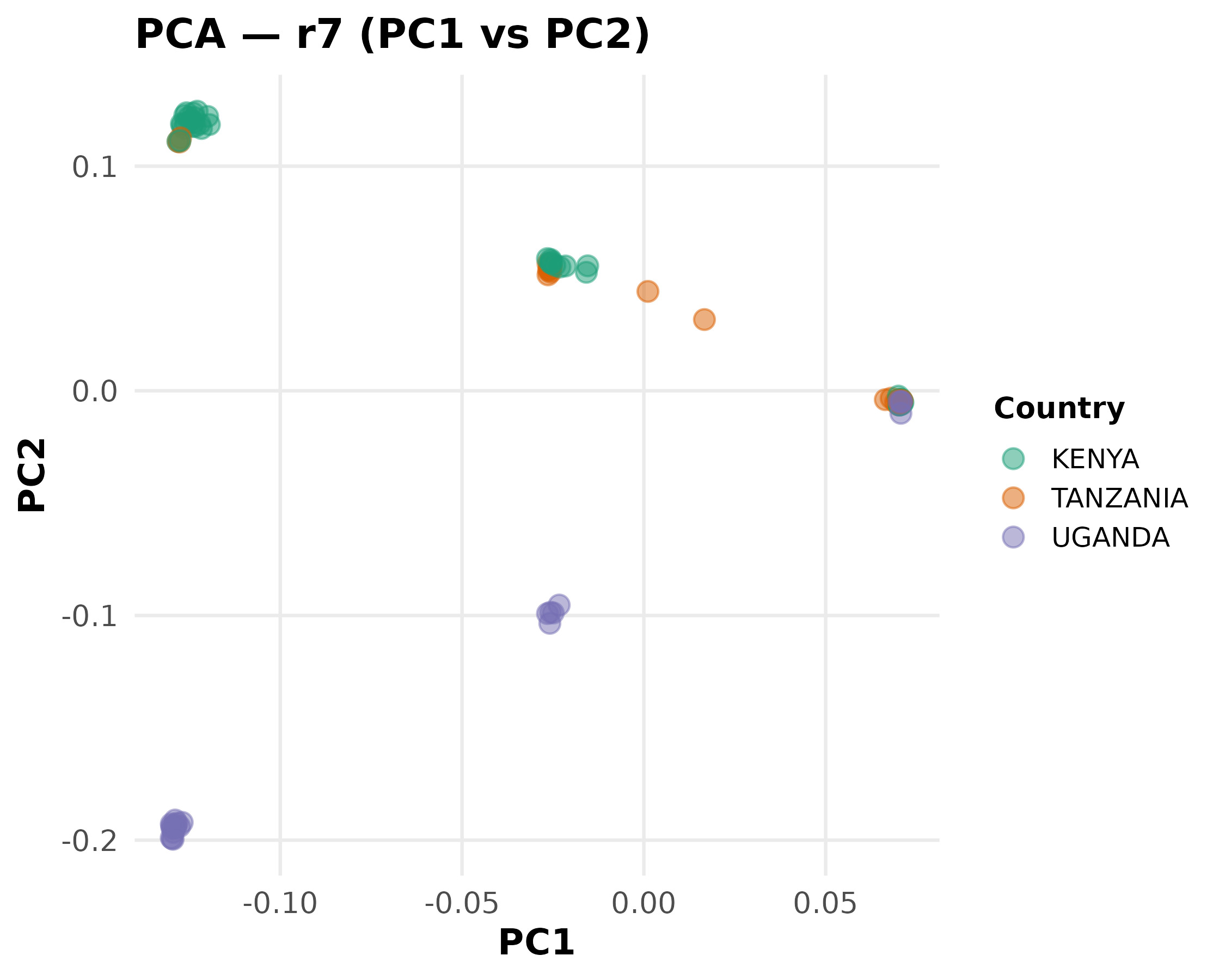

### Supplementary Figure 7

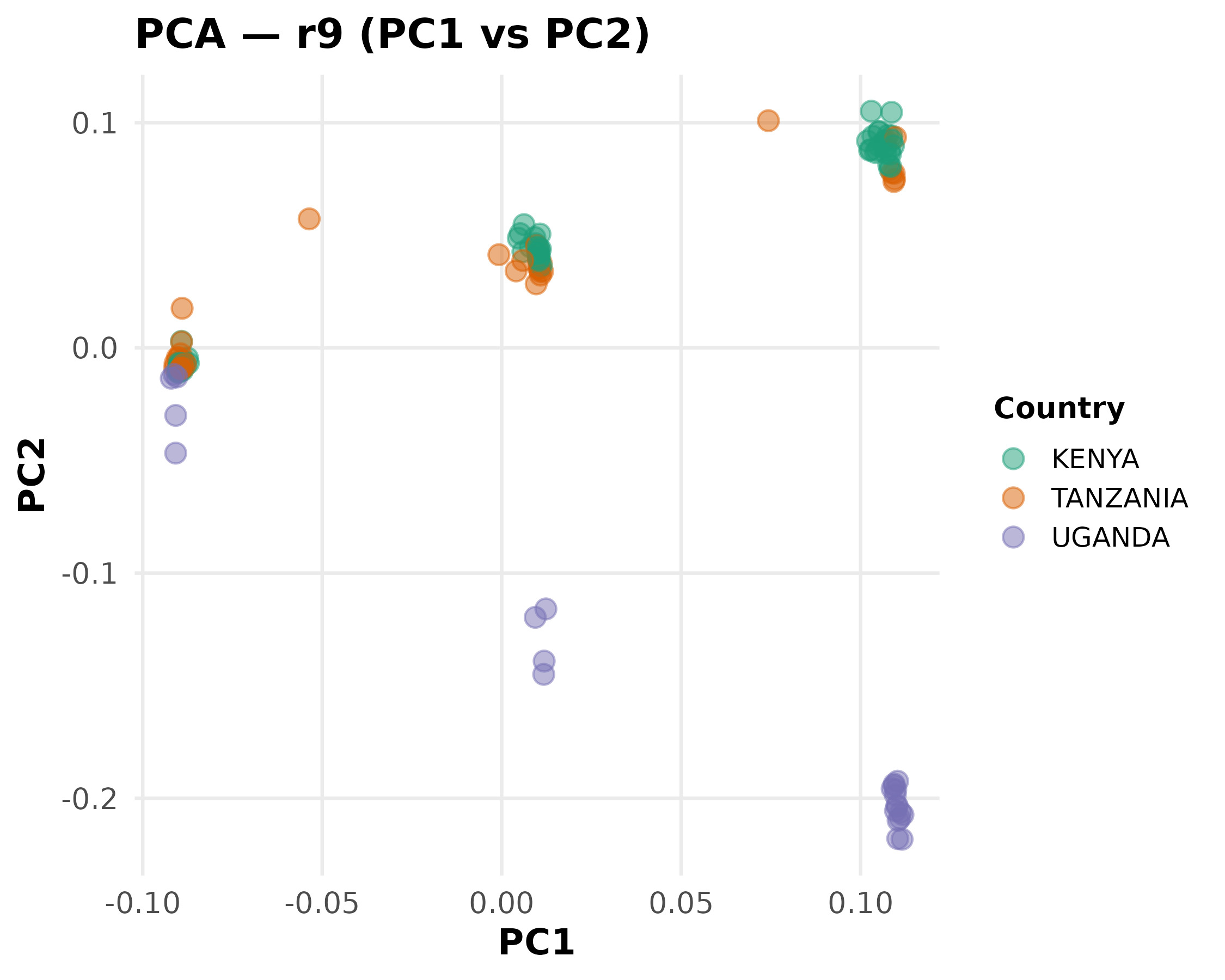

### Supplementary Figure 8

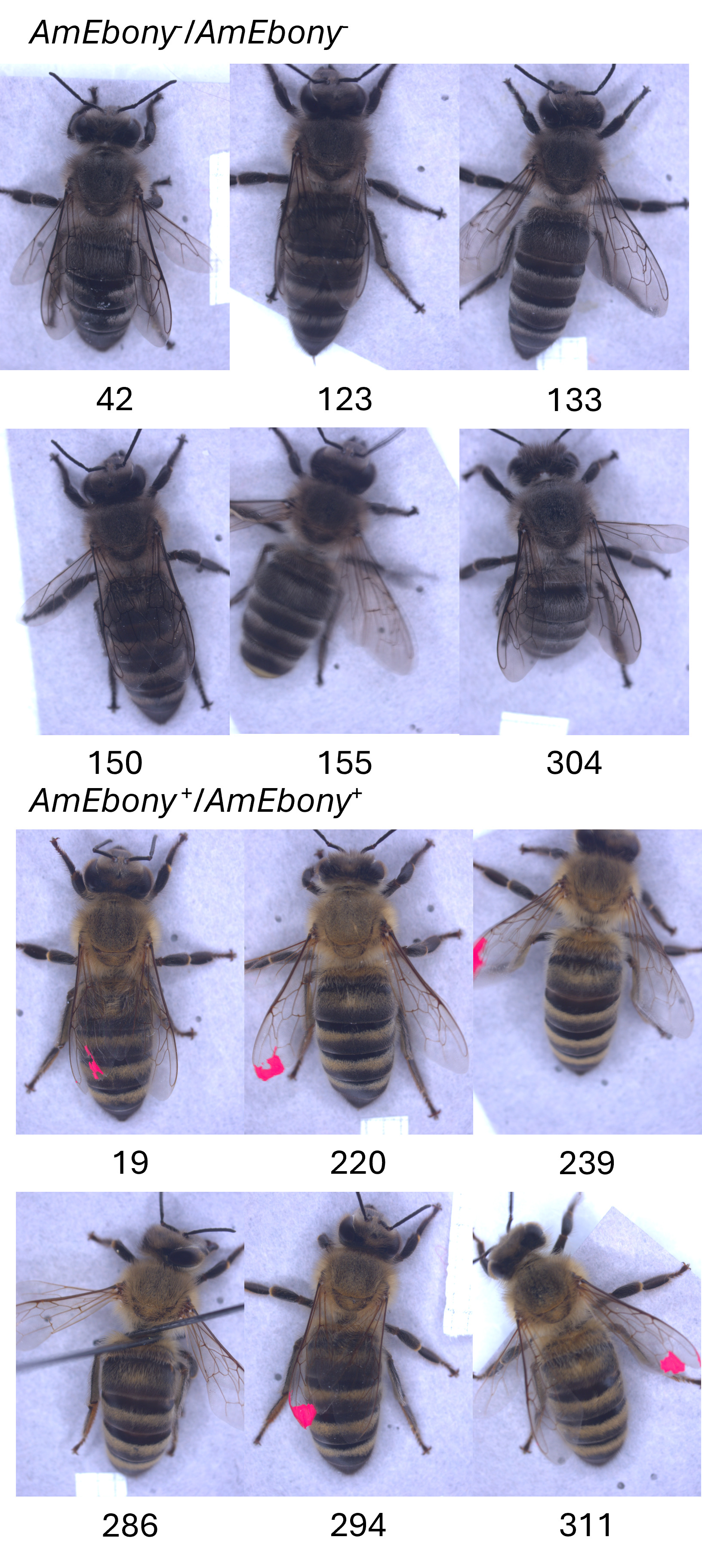
